# Parasites, infections and inoculation in synthetic minimal cells

**DOI:** 10.1101/2022.05.11.491559

**Authors:** Brock Cash, Nathaniel J. Gaut, Christopher Deich, Laura L. Johnson, Aaron E. Engelhart, Katarzyna P. Adamala

## Abstract

Synthetic minimal cells provide a controllable and engineerable model for an increasing amount of biological processes. While much simpler than any live natural cell, synthetic cells offer a chassis for investigating the foundations of key biological processes. Here we show a synthetic cell system describing host cells interacting with parasites and infections of varying severity. We demonstrate how the host can be engineered to resist infection, we investigate the metabolic cost of carrying resistance, and we show a simple inoculation system that immunizes the host against pathogens. Our work expands the synthetic cell engineering toolbox by demonstrating host pathogen interactions and mechanisms for acquiring immunity. This brings synthetic cell systems one step closer to providing a comprehensive model of complex, natural life.

## Introduction

Within the past several decades, the field of synthetic biology has exploded- allowing for construction of synthetic cells that resemble and model aspects of natural biological life. Modeling cellular pathways and behaviors in synthetic cells offers many advantages over live cells such as a more simplified and defined reaction environment, complete control over the proteomic and chemical makeup of the cell, the ability to compartmentalize interfering biological pathways and processes, as well as maintaining important biological systems that are absent in *in-vitro* reaction systems. ^1, 2^ These unique assets of synthetic cell technologies have already allowed for simulation of biological conditions and pathways within a life-like *in-vitro* bioreactor (synthetic cell) and brought a better understanding to many areas of study including: molecular crowding,^3^ lipid-protein dynamics,^4^ minimal metabolism,^5^ and many other critical aspects of biological life.^2^ These life-like synthetic cells with their unique assets are rapidly becoming better models and shedding more light into the mysterious inner-workings of cellular life.^6, 7^

These breakthroughs are not confined to single-cell processes; engineering interactions between populations of liposomes has proved a powerful technology as well. For example, design of a synthetic cellular system that mimics a predator-prey relationship has been developed utilizing light as the signal between these synthetic cell populations.^8^ Other complex work has been developed that allows for signal propagation throughout a synthetic cell droplet matrix, allowing fluorescence to spread in a spatial manner among the population.^9^ In addition to synthetic cell-synthetic cell interactions, there has been recent advancements in synthetic cell-live cell interfaces as well. For example, it has been shown that a synthetic cell designed to interface with a neural stem cell is able to induce neural differentiation as well as act as a generalized chassis capable of interfacing with other cell types.^10^ There have been countless advancements in the development of signal propagation and synthetic cells acting as interfaces, much of which is explored in the following reviews.^11–13^ In short, there are whole fields of research concerned with mimicking the interactions of cellular populations in the model system of synthetic cells.

Despite this, there has not been a study that uses synthetic cells to model infection, parasitism, or inoculation. Just as synthetic cells are commonly used to model complex biological processes in a simplified and controlled way, the potential exists to be able to model these complex inter-organism interactions. The importance of this can be seen in the groundbreaking study that successfully produced active phage particles within a cell-free protein expression platform.^14^ We aim to bring this concept to the macro-scale by designing synthetic cell populations that can actively infect one another and hijack a host cells’ genome, just as live viruses and parasites do. Being able to apply and engineer synthetic cells to model these disease behaviors will provide a valuable tool to the scientific community.

In this work, we exhibit our system for synthetic cell disease modeling. We engineered a host cell that could undergo infection from a second population of parasitic synthetic cells and investigate the resource competition, genome hijacking, and metabolic repurposing that occurs during such an infection. We also model a method that allows for rescue of the host after infection as well as a means of inoculation that will allow for the host to resist the infection of the disease-causing synthetic cells. In short, we present a novel disease and infection model chassis for synthetic cells. With this technology, synthetic cells can begin to be used as rudimentary models for disease, infection, and inoculation.

## Results and discussion

In this work, we use the words: host, parasite, infection, deadly, vaccine and inoculation – all in context of non-living, synthetic cell systems. By “deadly” infection we mean a marked decrease of translation in the population of synthetic cells. All experiments in this work were performed on synthetic cells that are not alive, do not self-replicate, and do not evolve. This is a model system for natural biological properties and processes.

### The parasite infection hijacks resources of the host

To test the concept of a synthetic cell system mimicking an infection, we started by constructing a model system for a parasite. We used the simplest concept of a parasite possible to reconstitute in this non-living biochemical system: a parasite that hijacks some of the resources of the host. In this case, the host metabolism is represented by expression of GFP protein, and the parasite is represented by mCherry protein expression. For this system, we utilized the previously demonstrated phenomenon of linear protein expression yield dependent on the DNA template concentration in TxTl. ^15, 16^ First we set out to find the optimal total plasmid concentration at which the ratio of the GFP and mCherry DNA corresponds to the ratio of observed fluorescence from those proteins. We arbitrarily assigned the GFP gene as the “host” and the mCherry gene as the “parasite.” Those names signify nothing special in this particular experiment, but will indicate host and parasite in later synthetic cell experiments. Here we use those names to make it easier to relate those proof-of-concept experiments to later synthetic cell infection experiments using the same plasmids.

We mixed GFP and mCherry plasmids, with both genes under the control of the T7 promoter, at varying plasmid ratios and total plasmid concentration in the sample- 1nM, 5nM, 10nM or 15nM (**Figure 2b**). The results followed the expected nearly linear relationship between the concentration of plasmid and the recorded fluorescence intensity, with the best match between fluorescence and plasmid concentration ratios observed at a total of 10nM plasmid. The observed relationship indicates that the plasmids, with the proteins being similar in size and amino acid composition (**Figure S1**), compete for the same pool of TxTl resources for their expression. Changing the relative concentration of the two plasmids results in changes in the expression efficiency roughly proportional to the ratio of the plasmids. This indicates it would be a good model to mimic parasite infection: the parasite genome (mCherry plasmid) will compete with the host genome (GFP plasmid).

**Figure 1.**
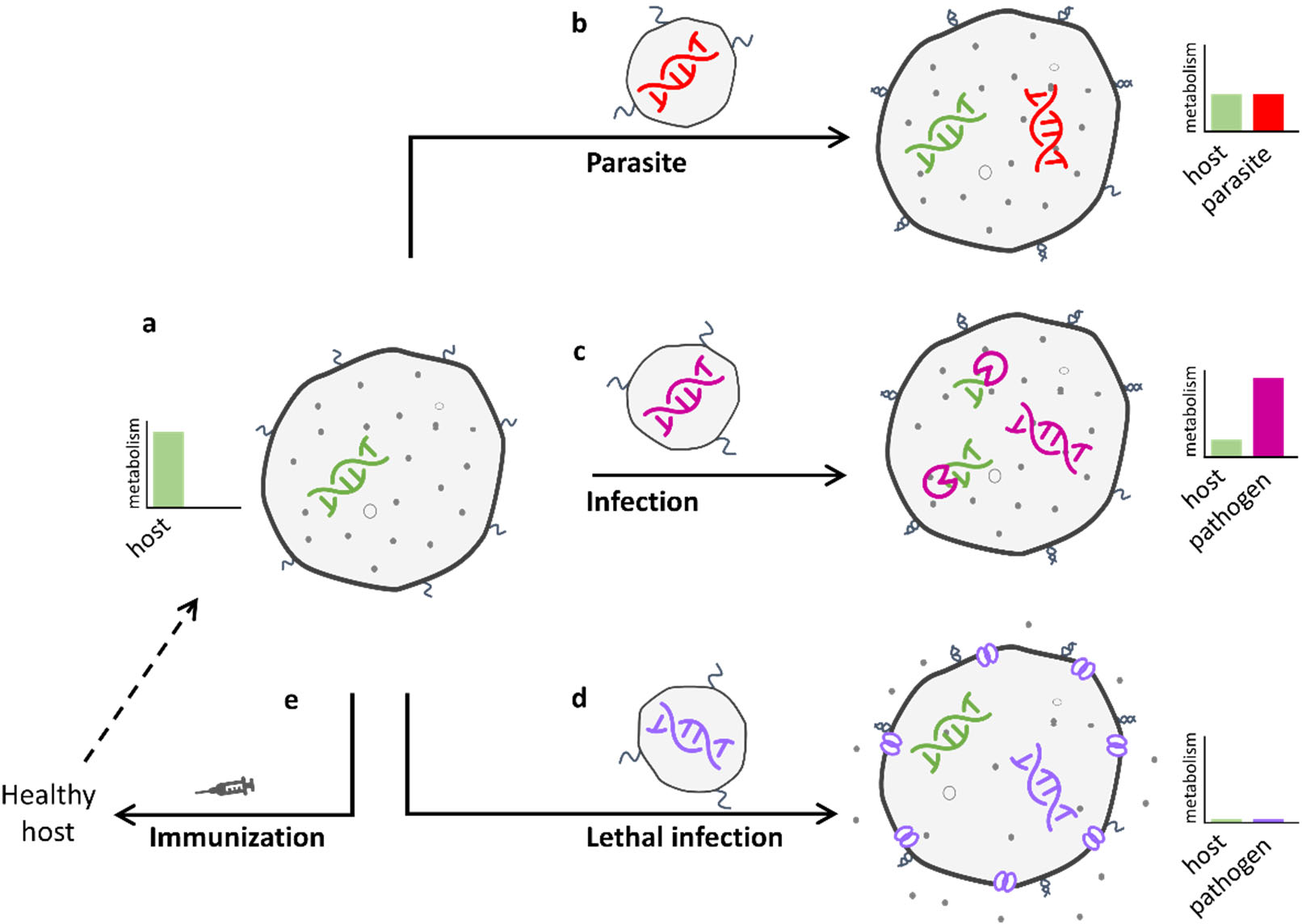
The parasite, infection, and inoculation system in synthetic minimal cells. **a**: **The “host”** is a synthetic cell expressing green fluorescent protein (GFP). On the surface of the outer membrane, the host carries ssDNA tags. The only proteins translated in a healthy host are the host’s own proteins from plasmids the host was created with. **b**: **The “parasite”** is a liposome filled with plasmid DNA encoding a gene for mCherry fluorescent protein, and the surface of the parasite liposome is decorated with ssDNA tag (complementary to the tag on the host surface). The “infection” starts with fusion of the host and parasite liposomes, mediated by the complementary DNA tags on their surface. After the infection, the host expresses both the original host gene (GFP) and the parasite gene (mCherry). **c**: **The “infection”** starts with “pathogen” liposomes carrying a gene for restriction enzyme MunI fusing with the host liposomes. The pathogen gene MunI is expressed by the host and the MunI restriction enzyme digests host DNA. This results in the native host genome being digested, turning most of the host protein expression from GFP to the pathogen’s encoded protein, MunI. **d**: **The “lethal infection”** is a fusion of the host liposome (with the GFP genome) with the pathogen liposome containing the gene encoding aHL, a membrane protein. After the infection, the host expresses aHL, which creates membrane pores, causing leakage of host nutrients and effectively shutting down the host’s metabolism. **e**: **“Immunization”** is incubation of the host with ssDNA containing a sequence compatible to the fusion tags on the surface of the host liposomes. The immunizing DNA creates a duplex with host DNA tags, preventing hybridization of other DNA oligos to those tags. This prevents fusion of any pathogenic liposomes to the host. This is a proof-of-concept figure, the graphs are for illustrative purpose, representing relative amounts of protein expressed from host and various pathogens. Real data for each system are on later figures.

**Figure 2.**
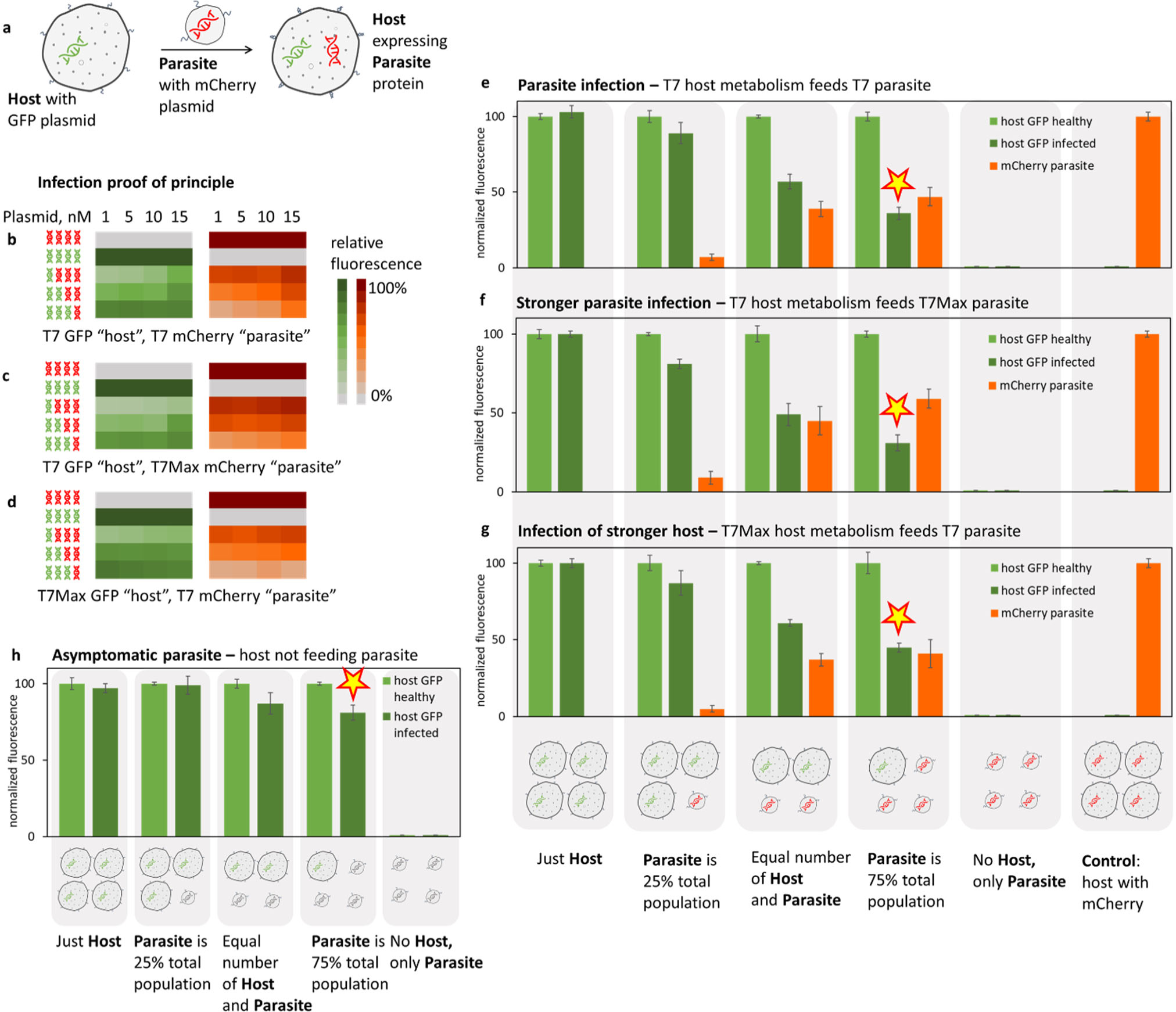
The parasite infection hijacks resources of the host. **a**: The host synthetic cell liposome contains a cell-free translation system and a plasmid encoding GFP. The surface of the host is decorated with ssDNA tags. The parasite liposome contains plasmid encoding mCherry, with a translation system but no RNA polymerase (so the parasite cannot express mCherry). The surface of the parasite is decorated with ssDNA tags complementary to the tags on the surface of the host. The DNA tags facilitate fusion of the host and parasite liposomes and the host expresses both its own genomic GFP and the parasitic genome containing mCherry. **b-d**: The proof of principle for the host (GFP) and parasite (mCherry) genomes competing for the same synthetic cell resources. Those experiments are done in solution TxTl, not in liposomes. Two plasmids, encoding GFP and mCherry, were mixed at the ratio indicated by the plasmid icons on the left of the heat map, with the sample’s total DNA concentration indicated in the row above the heat map. Fluorescence was measured in green (GFP) and red (mCherry) channel. Each plasmid was tested in two variants with promoters of different strengths: with T7 and with the stronger T7Max promoter. Individual data for all heat maps are on **Figures S2**, **S3** and **S4**. Representative individual time courses for the protein expression of the T7 host and parasite are on **Figure S5**, **S6** and **S7** and for the T7Max parasite on **Figure S8**, **S9** and **S10**. **e-g**: Infection of the host with the parasite. In each experiment, host liposomes are mixed with parasite liposomes. The fusion of the parasite to the host delivers parasitic mCherry DNA, that is being expressed alongside host GFP DNA. Different ratios of host to parasite were tested, as indicated by the icons below the bar graphs. The host genome under the T7 promoter was infected with a parasite with its genome under the T7 promoter (panel **e**) and with a parasite with its genome under the T7Max promoter (panel **f**), the host was also made with a T7Max genome and infected with the parasite with a T7 genome (panel **g**). Representative individual time courses for protein expression are on figure **S11**. Similar experiments at higher and lower total lipid concentrations are on **Figure S12**. Size exclusion purification traces showing vesicle stability post fusion are on **Figure S13**. **h**: Infection with incompatible parasite: the parasite carries T7 promoter, but noncoding DNA. Error bars indicate SEM, n=3. Fluorescence values on panels **e**-**h** are normalized to the blue fluorescence of a membrane dye, to normalize the results to concentration of host cells. All sample series labeled “host GFP infected” are experiments where the host and parasite had complementary DNA tags on the surface, enabling fusion of the liposomes, termed infection. Series labeled “host GFP healthy” show samples where the host and parasite were mixed at the ratios shown, but did not contain complementary fusion tags, making infection impossible. The stars on panels **e**, **f**, **g** and **h** indicates one particular infection condition that best illustrates the differences in the host and parasite expression strengths (panels **e**-**g**) and the benefits of host immunity (on panel **h**). Direct comparison of the host and parasite values signified by the star are on **Figure S14**. Data for infection with a parasite without any plasmid are shown on figure **S15**.

Next, we decided to test a case where either of the plasmids was under the control of a stronger promoter – thus mimicking a biased resource allocation scenario. To keep the resource competition aspect of the future “parasite” infection experiments, we kept both genes under the same RNA polymerase promoter (the T7 RNA polymerase, T7 RNAP), but varied its strength. We used a strong T7 RNAP promoter T7Max, which demonstrates significantly higher transcription yield, resulting in correspondingly higher translation yields, in TxTl. ^17^ We tested the reaction setup identical to the one described earlier: two plasmids at different ratios, tested at varying final total plasmid concentration. One of the plasmids was under the control of the strong T7Max promoter, while the other was under the canonical T7 promoter. In both test cases, the T7Max plasmid expression was strong enough that the fluorescence of the protein from a gene under T7Max did not significantly increase with the increased total concentration of the plasmid (**Figure 2c** and **2d**). It did not seem to matter if T7Max was driving GFP or mCherry, both proteins responded to the stronger promoter with a similar relative increase. This provided a promising model for infection with a “stronger” parasite (a parasite carrying gene under T7Max into a host under T7) and reverse, stronger host resisting weaker parasite (a host with genome under T7Max and a parasite carrying T7 gene).

Encouraged by those proof of principle experiments, we moved on to synthetic cell experiments.

The “infection,” in all cases, is the fusion of a host and parasite synthetic cell. The fusion is facilitated by DNA tags on the surface of the liposomes, using the DNA inducible fusion system described before. ^18^

Briefly, the synthetic cells were created with *E. coli* TxTl inside a membrane of POPC and cholesterol and the outer leaflet was decorated with single stranded DNA anchored to the membrane by cholesterol modification. If a pair of liposomes with complementary tags meet, the DNA tags hybridize, bringing the membranes together and initiating membrane fusion, which is followed by lumen mixing.

In our experiments, the host synthetic cell is the liposome containing TxTl with T7RNAP in addition to the genome of a plasmid-encoded GFP. Monitoring GFP fluorescence serves as a proxy for the “health” of the host metabolism. The incoming parasite contains a genome in the form of a plasmid encoding mCherry, the complete translation system, but no T7 RNA polymerase. This ensures that any observed changes in the host metabolism results from the genes brought in by the parasite and not from the fusion of the parasite and host liposomes, diluting the host cytoplasm.

The parasite infection experiments were carried out with the host membrane labeled with Marina Blue DHPE, a blue membrane lipid dye. This allowed us to normalize the observed host and parasite gene fluorescence to the blue fluorescence – normalizing the results to the total amount of host membrane fluorescence present in the sample. This accounts for the dilutions of the total volume of the sample after addition of parasite liposomes. The parasite membranes were not fluorescently labeled.

All experiments were accompanied by a control sample where only one population was present – the host, but with mCherry plasmid, to show the theoretical maximum possible mCherry expression levels if it was the only gene in the population.

In the first set of experiments, we mixed host and parasite with genes both under control of the regular T7 promoter – so both host and parasite genes were equally “strong.” We mixed the host with parasite so that the parasite was at 25%, 50%, or 75% of the total liposome population. In all cases, the host GFP fluorescence decreased in the presence of the parasite, with the corresponding increase of parasite mCherry fluorescence (**Figure 2e**). Samples with only the parasite and no host produced no fluorescence of either color, as expected. In all cases, the DNA-mediated liposome fusion is not 100% efficient;^18, 19^ therefore, the relative decrease in host fluorescence and increase of parasite fluorescence was not as large as the expected theoretical value based on the ratio of host to parasite. In other words, the infection was not 100% efficient. This serendipitous quality of DNA induced fusion system was, in this case, a beneficial feature of the system. As in the case of natural infection, the synthetic cell fusion mediated infection did not affect all hosts and not all parasites found a host.

Next, we investigated two cases where host and parasite were not evenly matched in strength of their genomes. In one case, the host genome was under control of the T7 promoter while the parasite genome was under the T7Max promoter (**Figure 2f**) and in the opposite scenario, the host genome was under T7Max and parasite genome was under the T7 promoter (**Figure 2g**). In the case of the weaker host attacked by a stronger parasite, the relative increase in parasitic mCherry fluorescence and corresponding decrease in host GFP fluorescence was much stronger than in the previous case of an evenly matched host and parasite. As expected, when the host was stronger than the parasite, the opposite was true: the mCherry fluorescence levels were lower and the relative decrease in GFP fluorescence was lower than in the evenly matched case. This is best illustrated by comparing a single data point under the same infection conditions: at 75% parasite ratio. Evenly matched host and parasite populations results in a parasite fluorescence slightly larger than the host. When the parasite is stronger, the parasite fluorescence is significantly higher than the host. And when the host is stronger, the parasite fluorescence is slightly lower than the host. These highlighted data points are marked with a yellow star on **Figure 2e**, **f,** and **g**.

We performed the control experiment of a host infected with an “asymptomatic” parasite – a parasite carrying a plasmid with a T7 promoter, but no protein coding sequence (we removed the mCherry cassette from the plasmid) (**Figure 2h**). The host GFP fluorescence decreases minimally at higher parasite levels, indicating some small amount of host resources are devoted to transcribing the short non-coding RNA from the parasite plasmid. This data also confirms that the main metabolic burden of the parasite infection on the host lies in utilizing translational, not transcriptional, resources. This observation falls in line with previous observations of translation being the more variable and limiting resource in cell-free systems. ^20^

Overall, those experiments provide a model for infection with a parasite, that hijacks metabolic resources of the host, with synthetic cell behaviors mimicking aspects of a natural host-parasite relationship.

### The infection destroys the host genome and hijacks host resources

The parasite described above was expressing a genome encoding mCherry using resources of the host, but the host maintained some of its metabolism and continued expressing GFP despite the infection. Next, we decided to investigate a system in which the infection results in a complete takeover of the host metabolism by the parasite via destruction of the host genome (**Figure 3a**).

**Figure 3.**
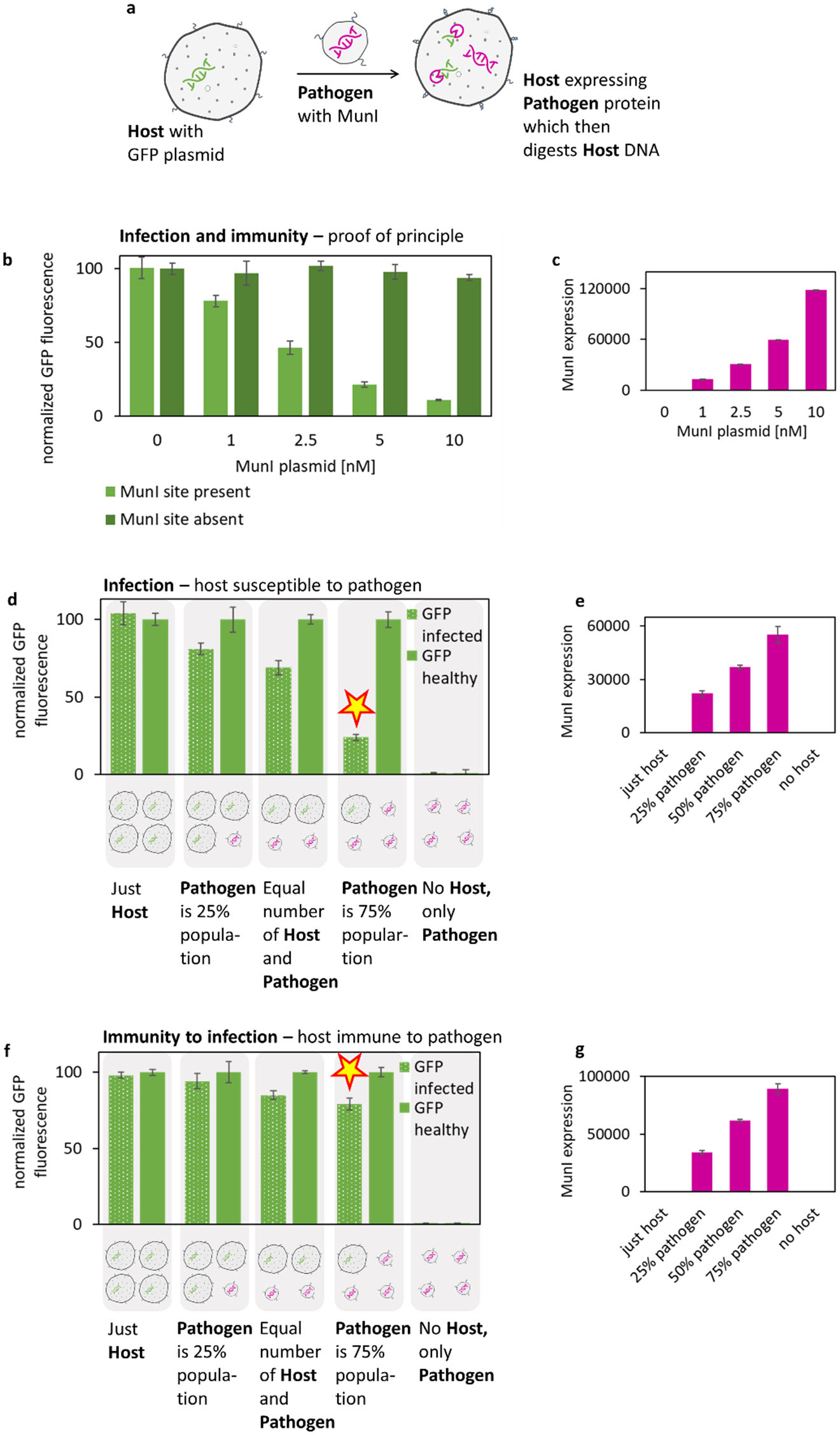
The infection destroys the host genome and hijacks host resources. **a**: The host cells contain plasmid encoding GFP, with two MunI restriction sites; the pathogen contains plasmid encoding the MunI restriction enzyme. After fusion of the host and pathogen, the host TxTl expresses MunI restriction enzyme, which then digests the host genome. **b**: The proof of principle experiments for the infection: GFP plasmid was prepared with and without MunI restriction sites. GFP plasmid was mixed with MunI plasmid and GFP expression was recorded. Those experiments were performed in solution TxTl, without liposomes. Proof of principle for the infection experiment, with MunI enzyme added externally instead of being expressed in TxTl, are on **Figure S16** and **S17**. Representative protein expression time courses are on **Figure S18**. **c**: MunI expression was monitored by Western blot analysis; the intensity of MunI band was quantified in the same samples as fluorescence results shown on panel **b** for the “MunI site absent” series. Representative western blot of MunI restriction enzyme is on **Figure S19**. **d**: GFP Fluorescence of the host mixed with pathogen liposomes at the ratio shown under the data graph. The host GFP plasmid has two MunI restriction sites. Representative protein expression time courses are on **Figure S20**. Data for infection experiments with incompatible fusion tags are on **Figure S21**. **e**: Expression of MunI was tracked via Western Blot analysis through the experiment for the samples shown in the series “GFP infected” on panel **d**. **f**: GFP fluorescence of the host mixed with pathogen liposomes at the ratio shown under the data graph. The host GFP has no MunI restriction sites, giving it immunity to the pathogen protein. Representative protein expression time courses are on **Figure S22**. **g**: Expression of MunI was tracked via Western Blot analysis through the experiment for the samples shown in the series “GFP infected” on panel **f**. Error bars on all panels indicate SEM, n=3. Fluorescence values on panels **d** and **f** are normalized to the blue fluorescence of the membrane dye used to normalize the results to the concentration of the host cells. Fluorescence on panel **b** is normalized to 0mM MunI plasmid for each sample series. The star on panels **d** and **f** highlights one particular condition best illustrating the differences in responses to the infection from a host susceptible to the pathogen (panel **d**) and immune to the pathogen (panel **f**). All the sample series labeled “GFP infected” are experiments where the host and pathogen had complementary DNA tags on the surface, enabling fusion of the liposomes, termed infection. The series labeled “GFP healthy” show samples where the host and pathogen were mixed at the ratios shown, but did not contain complementary fusion tags, making infection impossible.

To engineer such a scenario, we investigated a restriction enzyme digest of a plasmid in TxTl. In the proof of principle experiments, a MunI restriction enzyme was expressed in TxTl without liposomes. The MunI plasmid was mixed with GFP plasmid at different ratios. If the GFP plasmid contains no MunI restriction sites, the relatively small decrease in observed GFP fluorescence can be accounted for by the previously described competition between the two plasmids for TxTl resources. If the GFP plasmid has a MunI restriction site, the GFP fluorescence decreased significantly, and the decrease scaled proportionally to an increasing amount of the MunI-encoding plasmid (**Figure 3b**). The MunI expression was monitored via Western Blot and scaled proportionally with the plasmid concentration (**Figure 3c**).

Encouraged by those results, we set up an infection experiment with synthetic cells. The host carried GFP plasmid and the pathogen carried MunI plasmid. The concentration of GFP and MunI plasmids was set to 2mM, well below the concentration of plasmid saturation for TxTl. ^21^ We intended to remain under the total plasmid concentration at which host and parasite expression would be high enough to directly compete for limited resources (like in the above-described case of simple parasite), instead intending to mostly measure the effect of the parasite enzyme actively destroying the host genome.

Like in the earlier described parasite experiments, the host contained bacterial TxTl and T7 RNA polymerase, the pathogen carried only cell extract without T7 RNA polymerase and both host and pathogen were labeled with complementary DNA fusion tags. The host membrane was labeled with Marina Blue DHPE to normalize fluorescence results to the concentration of host liposomes.

The host and pathogen were mixed at varying ratios, with the pathogen being 25%, 50%, or 75% of the total population. The fluorescence of the “infected” host, where the host and pathogen contained complementary fusion tags, decreased with an increasing amount of pathogen. The GFP fluorescence from the “healthy” host, where the pathogen did not contain complementary fusion tags, did not decrease in the presence of any amount of pathogen. Samples with only the pathogen resulted in no measurable fluorescence (**Figure 3d**). The pathogenic MunI protein expression was quantified using Western Blot analysis, with data shown for the sample series of the infected host (**Figure 3e**).

Next, we investigated infection in the case of a host immune to the pathogen: the host GFP plasmid did not have a MunI restriction site (**Figure 3f**). The host fluorescence after infection decreased, but nowhere near as significantly as in the earlier case of a host with a MunI site in the genome. The MunI pathogen protein was expressed at levels comparable to earlier infection experiments (**Figure 3g**), so the metabolic load on the host was similar. The infected host fluorescence decreasing in those experiments is caused by some of the host resources being hijacked by the pathogen expressing MunI, but because the total plasmid concentration is much lower than in the earlier simple pathogen experiments, the host fluorescence decrease is less significant. This experiment also confirms that the decrease in fluorescence of the host with a MunI site (**Figure 3d**) was caused mainly by MunI digestion of the host genome, since the fluorescence of the host without a MunI site on the genome does not respond to infection as much (**Figure 3f**). The healthy host in those experiments was again not affected by addition of a pathogen without complementary fusion tags.

The yellow star on **Figure 3** indicates experiments directly comparable as examples of host immunity (conveyed by the lack of a restriction site): on panel **d,** the host fluorescence in the case that 75% of the liposome population is composed of the pathogen is approximately a quarter of what the fluorescence of the immune host is in the case of the same pathogen to host ratio shown on panel **f**.

### The host is susceptible to “deadly” infection, but the host can be rescued

The above parasite and pathogen experiments demonstrated that synthetic cells can be used as a model for infection where a pathogen hijacks resources of the host. Next, we decided to investigate a scenario in which the infection causes the metabolism of the host to cease functioning after expression of the pathogenic protein. In this case, the pathogen is comparable to infection with a bacterium expressing a deadly toxin – minus the replication aspect of the host or parasite, since those synthetic cells do not replicate. (**Figure 4a**)

**Figure 4.**
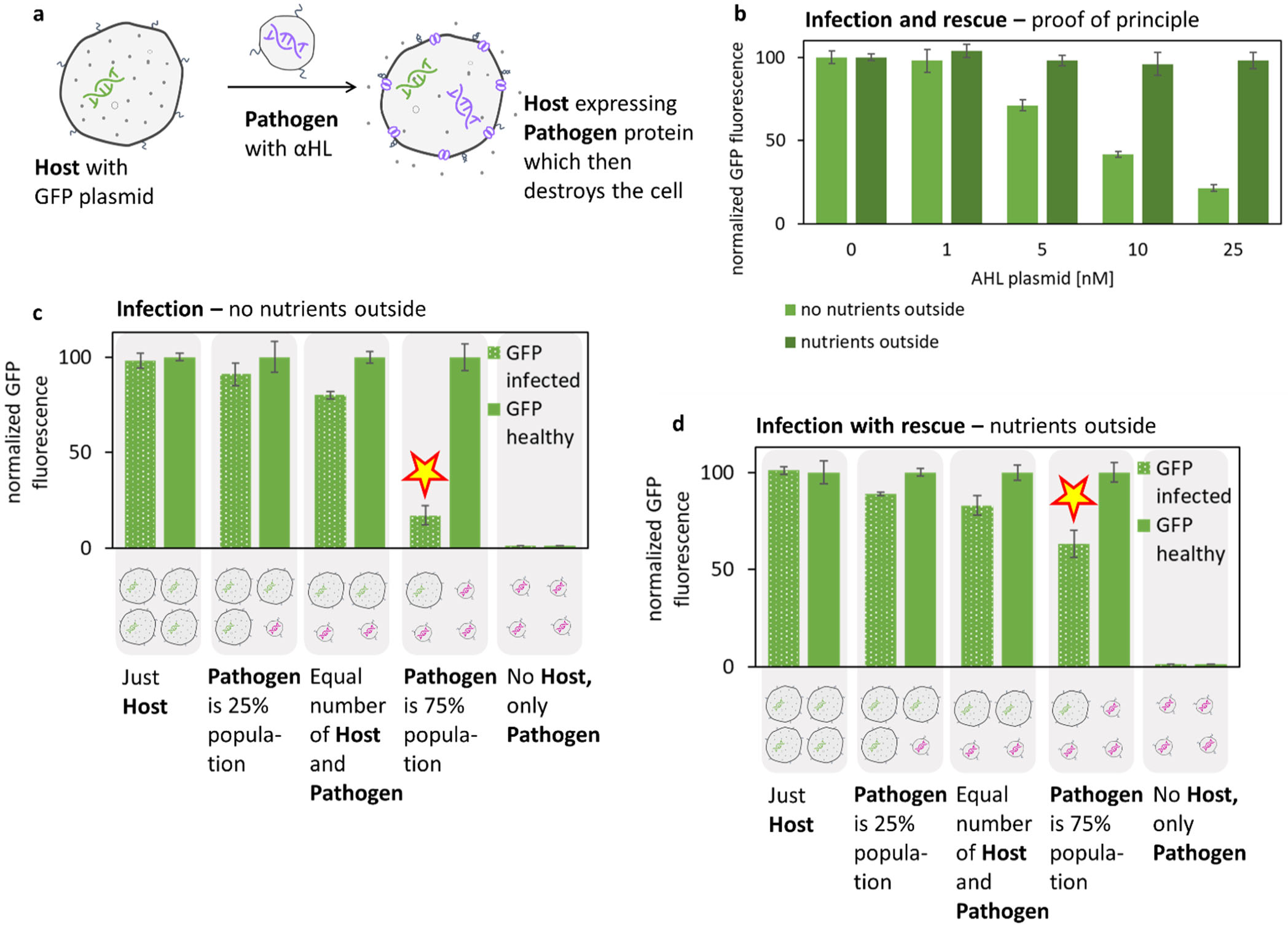
The host is susceptible to “deadly” infection, but the host can be rescued. **a**: The host contains GFP plasmid, the pathogen contains plasmid encoding the alpha hemolysin (aHL) membrane channel. After the infection, the host expresses aHL, which creates pores in the membrane, causing leakage of nutrients, and resulting in a decrease of metabolic activity of the host. **b**: Proof of principle for a host expressing aHL (no infection, host was created with both 5nM GFP and the indicated amount of aHL plasmid). The host was incubated in buffer with or without nutrients (all components of the energy, salt, and amino acid mix used to prepare host cell-free translation system). The optimum rescue nutrient concentration was established experimentally, data on **Figure S23**. **c**: Infection of the host with the pathogen, with varying amount of pathogen per host cell. The leakage of small molecules from the “healthy” and “infected” cells was directly measured using small molecule dye, data on Figures S24 and S25. **d**: Experimenta similar to those shown on panel c, but the outside buffer contains small molecule nutrients (similar to “nutrients outside” experimental conditions shown on panel **b**). Error bars on all panels indicate SEM, n=3. The fluorescence values on panels **b**, **c** and **d** are normalized to the blue fluorescence of the membrane dye used to normalize the signal to the concentration of the host cells. The stars on panels **c** and **d** highlights one particular condition best illustrating the differences in response to the infection from a host without outside nutrients (panel **c**) and immune to the pathogen due to the presence of outside nutrients (panel **d**). For panels **c** and **d** the series “GFP infected” is the host fused with the pathogen carrying aHL protein and “GFP healthy” is the host fused with the pathogen carrying no plasmid.

To engineer this system, we chose alpha hemolysin (aHL), a natural toxin and membrane protein widely used in synthetic cell engineering to create nonspecific membrane pores. ^22, 23^ We appreciate the irony of this choice, as aHL is naturally expressed by *Staphylococcus aureus* as a toxin and the protein was later appropriated to engineer membrane transport and prolong synthetic cell metabolism.^24^ Here we revert to using aHL for its natural role as a harmful toxin produced by an infection. Another notable synthetic cell engineering related example of using aHL’s toxicity to cause natural cell death comes from the application of synthetic cells as a cancer therapy technology. ^25^

As a proof of principle, we established that expressing a high concentration of aHL in synthetic cell vesicles results in leakage of cell content and a significant decrease in expression of GFP inside the cell. This effect can be reversed if the outside of the liposomes contains all the small molecules necessary for TxTl (**Figure 4b**).

In our host pathogen model system, the host contains GFP-encoding plasmid and the pathogen contains aHL-encoding plasmid. All the infection experiments’ design is similar to the parasite and pathogen scenarios described above, with the pathogen carrying no T7 RNAP and both the host and pathogen decorated with DNA fusion tags. Interestingly, we observed that at lower pathogen concentrations (the pathogen being 25% or 50% of the total population) the host fluorescence decreases only slightly. However, increasing the pathogen concentration to 75% results in a significant decrease in the host GFP fluorescence (**Figure 4c**). We speculate this result might be caused by the relatively slow leakage of nutrients from synthetic cells at a lower aHL concentration. The GFP fluorescence was measured only after 12 hours of incubation and GFP is a very stable protein. Therefore, it is possible that in the case of lower aHL concentrations (lower pathogen amount), the aHL-caused leakage of nutrients was slow enough that the host cells accumulated significant amounts of GFP before the effects of the aHL-induced leakage became crippling to the metabolism.

To decouple the results of simple competition for resources between the host and parasite genomes, we performed “infection with rescue” experiments in which the outside buffer contained all the small molecules that compose the TxTl energy, amino acid, and salt mixes. This was setup similar to the “nutrients outside” condition tested on **Figure 4b**. The decrease in host fluorescence in those experiments was similar to the earlier “infection without rescue” samples with the 25% and 50% levels of the pathogen. The decrease of host fluorescence at 75% of pathogen was significantly smaller than the decrease observed in samples without rescue (**Figure 4d**). The big difference in results for the experiment with 75% of the parasite, with data points on **Figure 4** panels **c** and **d** highlighted with a star, seem to support our hypothesis that higher aHL concentrations cause harm early in the reaction. At lower pathogen concentration the leakage is slow enough that the host GFP accumulates (as seen both with and without rescue nutrients), so the observed GFP fluorescence decrease after infection is mostly due to the competition for resources. At higher pathogen concentrations, the GFP fluorescence in samples with external nutrients decreases mostly due to resource competition.

Those experiments demonstrate that we can engineer synthetic cells to model infection resulting in two different outcomes: the metabolic penalty of the infection itself (decreased GFP due to resource competition) and a more “deadly” infection resulting from abolishing the hosts’ protein expression. Encouraged by this unexpected observation, we decided to investigate a system in which synthetic cells could actively resist a pathogenic infection, and investigate the metabolic penalty of such a response.

### Host cells defend against infection

We expanded on the parasite system presented on Figure 2. In addition to GFP, we added MunI- containing plasmid to the host. The parasite contained mCherry, as in previous experiments. The mCherry plasmid contains a MunI restriction site (**Figure 5a**).

**Figure 5.**
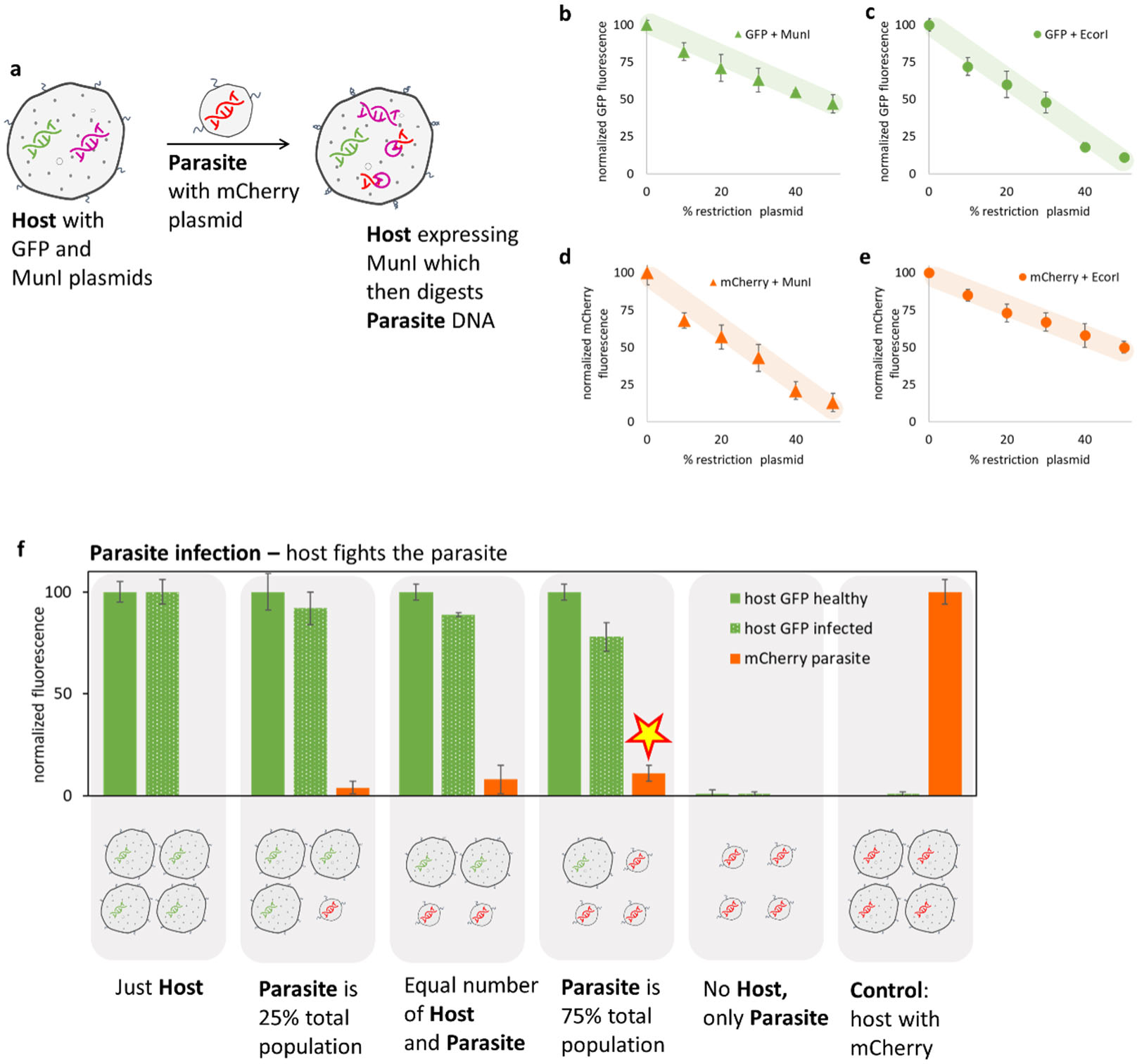
Host cells defend against infection. **a**: The host cells contain two plasmids: one encoding GFP (without a MunI restriction site) and the other encoding MunI restriction enzyme (the “parasite resistance” gene). The parasite contains mCherry plasmid, in a system similar to experiments shown on Figure 2. The infection is mediated by fusion via DNA tags on the surface of the host and parasite. After the infection, the mCherry plasmid introduced by the parasite is digested by the host’s MunI restriction enzyme. **b**-**e**: Measuring the metabolic cost of expressing the parasite resistance. Experiments are in a bulk TxTl system, not encapsulated in vesicles. In each experiment, fluorescent protein plasmid (GFP or mCherry) was mixed with restriction enzyme plasmid (MunI or EcorI) at the ratio indicated on the x-axis, for the total plasmid concentration of 10mM. GFP has an EcorI restriction site, but not a MunI site and mCherry has a MunI, but not an EcorI restriction site. The trendline is a visual guide, not a data fit. Error bars indicate SEM, n=3. Control experiments with purified restriction enzymes, instead of expressing them from a plasmid, are on **Figure S26**. The stability of liposomes with an increased amount of parasite plasmid was also measured, data on figure **S27**. **f**: Infection of the host with an increasing amount of parasite. “Host GFP healthy” is the expression of GFP from host cells fused with the parasite without any plasmid. “Host GFP infected” is the host fused with the parasite carrying mCherry. The red fluorescence is mCherry fluorescence from the parasite fused with the host (Only data from the infected host are available. The healthy host samples did not carry mCherry plasmid). Error bars on all panels indicate SEM, n=3. Fluorescence values are normalized to the blue fluorescence of the membrane dye used to normalize the results to the concentration of the host cells. The star indicates expression of the parasite protein under particular infection conditions, directly comparable to the data point highlighted with a star on Figure 2e – an experiment under the same conditions, but with the host lacking the MunI resistance gene.

First, we performed proof of principle experiments to assess the metabolic load of expressing MunI on the host. In those unencapsulated TxTl experiments, we monitored expression of GFP and expression of mCherry, each mixed with an increasing amount of either MunI or EcorI plasmid. All genes were under the control of the same T7 promoter. The GFP gene had an EcorI, but not a MunI, restriction site and the mCherry gene had a MunI, but not an EcorI, restriction site. When a restriction enzyme plasmid was mixed with a fluorescent protein plasmid without a recognition site for that restriction enzyme, the fluorescent protein expression decreased proportionally to the increase of restriction enzyme plasmid – indicating competition between the two plasmids for TxTl resources, as demonstrated earlier for GFP and mCherry on **Figure 2**. Those non-cutting pairs were GFP with MunI (**Figure 5b**) and mCherry with EcorI (**Figure 5e**). When a fluorescent protein was paired with a restriction enzyme capable of cutting the gene for that fluorescent protein, the fluorescence decreased significantly to nearly background levels at higher restriction enzyme plasmid concentrations. Those cutting pairs were GFP with EcorI (**Figure 5c**) and mCherry with MunI (**Figure 5d**). This indicated a decrease in fluorescence significantly beyond the simple competition for resources, and similar to the infection scenario demonstrated on **Figure 3**.

MunI expression in the presence of GFP decreases GFP expression. If we use GFP as a proxy for host “baseline” metabolism, then MunI can be seen as metabolic load on the host.

We proceed to the infection experiments, mixing host (with GFP and MunI plasmids) with parasite (with mCherry plasmid). Like previously, we investigated scenarios with an increasing amount of parasite in the population (**Figure 5f**). The positive control of a host expressing mCherry only (with EcorI instead of MunI to balance the metabolic load) created a benchmark for the theoretical maximum amount of pathogen protein that can be expressed in the host (**Figure 5f**).

As the amount of parasite increased, the host’s GFP fluorescence decreased, but the decrease in infected host fluorescence was significantly smaller than the decrease of GFP expression observed in earlier experiments with the same parasite. GFP and parasite mCherry fluorescence of this infected host expressing MunI (data point labeled with star on **Figure 5f**) can be directly compared with similar infection hosts without MunI (data point labeled with star on **Figure 2e**). In the host expressing MunI, the host GFP started with an absolute lower level while healthy (because some resources are diverted to expressing MunI) but after the infection, the host fluorescence decrease is much less significant and the host makes much less parasite mCherry protein than in the case of the host not expressing MunI. This experiment demonstrates the benefit of the host spending some resources on defense against parasites. This system could be seen as a very simple biochemical model for organisms expressing proteins to defend against future infections or even harmful environmental factors. The host originally is less robust (if we take GFP expression as a benchmark for host health), but in case of an infection, the additional resources the host spent on expressing defensive proteins pay off, providing protection against the infecting parasite. Therefore, in this simple nonliving synthetic cell, we reconstituted an example of fundamental biological arms race.

### Inoculation provides defense against infections

In the above-described example, we demonstrated the host defending against the influence of a pathogen’s genome by expressing an extra protein. Next, we asked if it is possible to model a scenario where some intervention makes a previously susceptible host immune to the infection in the first place. We called it an inoculation model. While recognizing this is far from a perfect analogy, the aspect we focused on is that the host synthetic cell does not expend any resources on generating immunity.

In our immunization model, we designed a treatment that turns the host synthetic cell that is susceptible to infection into cells that cannot fuse with a pathogen. The synthetic cell fusion system used throughout this paper relies on single stranded DNA tags to induce liposome membrane fusion. To inoculate against fusion with a pathogen, we incubated the cells with DNA oligonucleotide complementary to the fusion oligo on the surface of the cells (**Figure 6a**).

**Figure 6.**
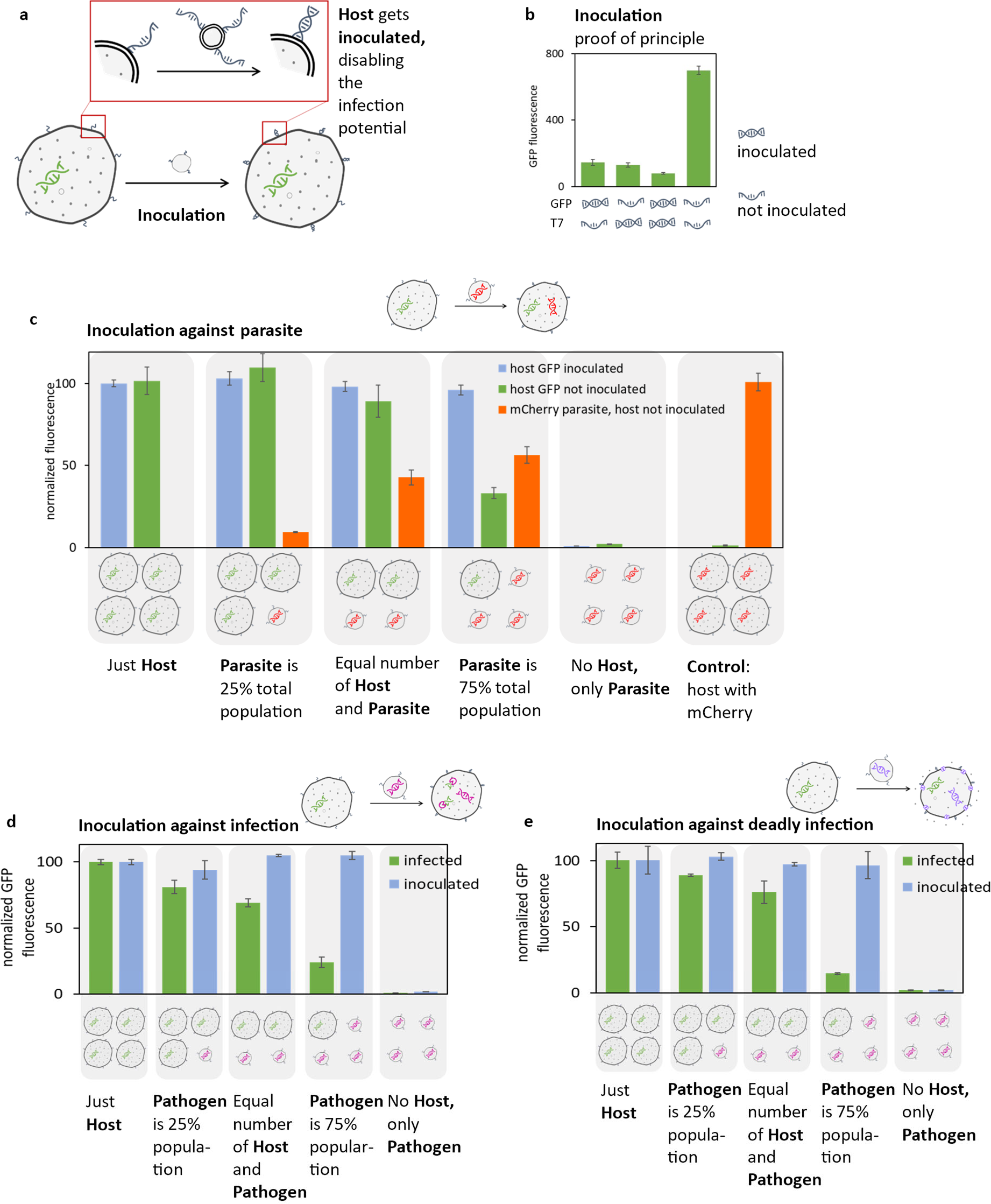
Inoculation provides defense against infections. **a**: The host contains GFP plasmid. The pathogen in those experiments is one of the three infection schemes demonstrated in this paper: parasite (as shown on Figure 2), pathogen with Munl restriction enzyme digesting the host’s GFP genome (as shown on Figure 3), or a pathogen carrying the membrane channel aHL causing leakage of nutrients from the host (as shown on Figure 4). In each case, the host and pathogen contain complementary DNA fusion tags. Before introducing the pathogen, the host is “inoculated”: incubated with excess ssDNA oligos complementary to the fusion DNA tags on the surface of the host (tags identical to those on the pathogen, minus the cholesterol moiety used to anchor the tag to the membrane of the pathogen). The inoculation causes the fusion tags on the surface of the host to become double stranded, thus making the fusion with the pathogen impossible. **b**: The proof of principle for inoculation scheme. The population of liposomes carrying GFP under the control of the T7 RNA promoter, but without T7 RNA polymerase, is fused with liposomes containing T7 RNA polymerase. The GFP protein can be expressed only if the two populations fuse. One or both of the mixing partners were “inoculated” by incubation with a DNA tag compatible to the fusion tags on the surface of the liposome, rendering the fusion tags double stranded and thus incompatible with fusion. A representative time course for protein expression is on **Figure S28**. Control experiments without any fusion tags are on **Figure S29**, and controls with incompatible fusion tags is on **Figure S30**. The inoculation oligo concentration was established experimentally, data on **Figure S31.** **c**: Parasite infection experiments. A host carrying GFP was fused with a parasite carrying mCherry, as described on Figure 2. The host and parasite were mixed with a varying ratio of host to parasite; host cells were either inoculated (with double stranded DNA fusion tags not compatible with fusion) or not inoculated (thus susceptible to fusion with parasite). mCherry fluorescence is reported only for samples with the host not inoculated, because there was no measurable mCherry fluorescence in the inoculated host samples. A reversed inoculation was tested by inoculating the parasite instead of the host, data on **Figure S32**. **d**: Infection experiments with the host genome susceptible to the pathogen carrying MunI restriction enzyme, as described on Figure 3. Infected data series represents a host without inoculation and the inoculated data series represents samples with the host incubated with inoculation oligos, thus not susceptible to fusion with the pathogen. **e**: Infection experiments with the pathogen carrying aHL gene, resulting in aHL channels leaking nutrients out of the host cells after infection, as described on Figure 4. Infected data series represents the host without inoculation and the inoculated data series represents samples with the host incubated with inoculation oligos, thus not susceptible to fusion with the pathogen. Error bars on all panels indicate SEM, n=3. Fluorescence values on panels **c**, **d** and **e** are normalized to the blue fluorescence of the membrane dye used to normalize the results to the concentration of the host cells. Some data shown on this figure are experiments with setup identical to the experiments shown on earlier figures (as indicated in each caption). Those experiments were repeated for this figure, to produce data using the same batch of TxTl for directly comparing the results, eliminating batch to batch variability of TxTl preparations.

First, we needed to confirm that the inoculation by incubation with complementary oligo works to abolish fusion. In the proof of principle experiments, two populations of liposomes were prepared: one population had TxTl with a plasmid encoding GFP under the control of the T7 promoter and the other population had TxTl and T7 RNA polymerase, but no plasmid. After fusion of those two populations, the GFP plasmid mixed with T7 RNA polymerase and induced GFP expression. If the two populations had single stranded and complementary fusion tags, fusion occurred and GFP fluorescence was detected. If one or both populations were incubated with oligonucleotide complementary to the fusion tag, the liposome fusion did not occur and GFP was not expressed (**Figure 6b**).

Having established the inoculation system, we proceeded to introduce inoculation to protect host synthetic cells in three of the infection scenarios described earlier in this work. First, we used the parasite scenario described on **Figure 2**. The host cells, containing a GFP genome, were inoculated with an oligo complementary to the fusion tag on the surface of the host and the host was mixed with the mCherry parasite. We mixed the host with the parasite at different ratios. Under all infection conditions, the parasite mCherry fluorescence was undetectable if the host was inoculated and the GFP fluorescence of the inoculated host did not decrease. The control experiments with the non-inoculated host showed a decrease in the host GFP expression and an increase in the parasite mCherry signal, as expected (**Figure 6c**).

Next, we applied the inoculation system to the case of a host infected with a pathogen carrying a genome encoding MunI restriction enzyme capable of digesting the hosts’ GFP genome, like in experiments shown on **Figure 3**. The inoculated host did not show a decrease in GFP fluorescence after mixing with increasing amounts of the pathogen, while the non-inoculated host was infected and the host GFP fluorescence decreased proportionally to the increasing amount of pathogen (**Figure 6d**).

Inoculation was also effective against the aHL pathogen. The inoculated hosts showed no decrease in fluorescence upon mixing with the pathogen, while the non-inoculated host’s GFP fluorescence decreased significantly with infection (**Figure 6e**).

In all cases, this simple inoculation procedure, incubating the host with a DNA oligo complementary to the fusion oligo on the host’s surface, was sufficient to abolish the ability of the pathogenic synthetic cells to infect the host cells. Therefore, we demonstrated a simple physicochemical model of intervention that protects host cells from infection.

## Conclusions

In this paper, we demonstrate methods for using synthetic minimal cells to model parasite and pathogenic infections. The motivation behind our work was two-fold. We wanted to engineer complex, life-like behavior into synthetic cells, furthering the goal of engineering synthetic cells that more closely resemble natural live cells. We also wanted to take advantage of the simple and controllable synthetic cell system to model basic properties of infection, inoculation, and resistance processes. While the synthetic cells do not evolve or self-replicate, this system allows for studying infection dynamics in a simple and easy to analyze model. We hope the principles of infection described here will help to create practical and applicable models for different pathogens and infection mitigation.

While the host infection dynamics presented here utilize bacterial based synthetic cells, this system is not inherently limited to bacterial TxTl. It would even be possible to imagine using the system presented here to model infection dynamics in populations resembling natural ecosystems, taking advantage of the simplicity inherent to synthetic cells to fully control and study the interactions between different members of large populations.

## Materials and methods

### Cell-free protein expression

All TxTl experiments were performed using *E coli* cell extract prepared according to the published protocol.^26^ We used Rosetta 2 (DE3) *E coli* strain to prepare all extracts.

For cell extract preparation, cells were grown in 2xYTPG media at 30°C to OD of 0.5. For each TxTl reaction, the final concentration of reagents was: from energy mix: 500mM HEPES pH 8, 15mM ATP and GTP, 9mM CTP and UTP, 2 mg/mL of E. coli tRNA mixture, 0.68 mM folinic acid, 3.3 mM nicotinamide adenine dinucleotide (NAD), 2.6 mM coenzyme-A (CoA), 15 mM spermidine, 40 mM sodium oxalate, 7.5mM cAMP, 300mM 3-PGA; from amino acid mix: 2mM each of Alanine, Arginine, Asparagine, Aspartic acid, Cysteine, Glutamic acid, Glutamine, Glycine, Histidine, Isoleucine, Leucine, Lysine, Methionine, Phenylalanine, Proline, Serine, Threonine, Tryptophan, Tyrosine and Valine; from salt mix: 130mM potassium glutamate, 10 mM ammonium acetate, 10 mM magnesium glutamate.^27^

Due to the batch-to-batch variability of TxTl preparation yields, we performed all the directly comparable experiments (experiments shown on the same figure) using the same batch of the extract.

### Liposome experiments

The synthetic cell liposomes were created using an inverted emulsion protocol described previously. ^28^

Liposomes were formed from dioleoylphosphatidylcholine (DOPC), dioleoyl-sn-glycero-3- phosphoethanolamine (DOPE), and cholesterol at a molar ratio of 3:1:1 ^29^. Thin lipid films were prepared by mixing all lipids in chloroform (in dark amber vials) and evaporating the solvent overnight. 500 µL of mineral oil was added to each thin film vial, incubated at 60°C for 10 minutes, vortexed for 10 minutes, incubated at 60°C for 3 hours, then sonicated in a bath sonicator at 60°C for 30 minutes. The mineral oil lipid samples were cooled down to 4°C. 30µL of internal liposome solution was added to each mineral oil sample, vortexed for 30 seconds, and equilibrated for 10 minutes at 4°C. The mineral oil liposome solution was carefully added on top of 250 µL of centrifuge buffer (100 mM HEPES + 200 mM glucose, pH 8). Samples were centrifuged at 18,000 rcf at 4 °C for 15 minutes.

As much of the mineral oil layer as possible was removed. Using a fresh pipette tip, the liposome “pellet” was aspirated from the bottom of the tube. The liposomes were resuspended in 250 µL wash buffer (100 mM HEPES + 250 mM glucose, pH 8), and centrifuged at 12,000 rcf at 4 °C for 5 minutes. Residual mineral oil was removed from the top of the solution and the liposomes were transferred to a fresh tube.

### Fusion (infection) experiments

The liposome fusion assay was performed as previously described.^19, 30, 31^ Briefly, the DNA fusion tags in 100 mM HEPES pH 8.0 were added to a solution of pre-formed liposomes. Liposomes were mixed immediately after the addition of DNA, and the samples were tumbled for 30 minutes to allow for DNA incorporation into liposome membranes. The DNA fusion tags sequence were 5’ GTCTAGCGTCTCACCAG/3CholTEG/ and a matching reverse complement. The fusion tags were labeled with CholTEG, a Cholesterol-TEG (15 atom triethylene glycol spacer) modification.

To account for the varying amount of host synthetic cell liposomes in samples with a varying ratio of host to pathogen, we labeled the host cells with Marina Blue DHPE (Marina Blue 1,2-Dihexadecanoyl-sn-Glycero-3-Phosphoethanolamine) at 0.1mol% in the lipid membrane

The fluorescence measurements were done on a SpectraMax plate reader recording GFP fluorescence (excitation/emission wavelengths 488/509 nm), mCherry fluorescence (excitation/emission wavelengths 587/610 nm) and Marina Blue fluorescence (excitation/emission wavelengths 365/460 nm) from the same sample, with PMT set to “medium” for all measurements, and with 12 reads per well.

## Acknowledgments

We are grateful to Thrasyvoulos Karydis for discussions on restriction enzyme activity in cell-free systems, which gave rise to ideas fundamental to conceiving this project. This work was supported by NASA award 80NSSC18K1139 Center for the Origin of Life - Translation, Evolution And Mutualism, NSF award 1844313 RoL: RAISE: DESYN-C3: Engineering multi-compartmentalised synthetic minimal cells, NSF award 1840301 RoL:FELS:RAISE Building and Modeling Synthetic Bacterial Cells, NSF award 2123465 Synthetic P-bodies: Coupling gene expression and ribonucleoprotein granules in synthetic cell vesicles for sensing and response, generous gift from Jeremy Wertheimer, and Hackett Royalty Fund award.

## Supplementary information

**Figure S1.**
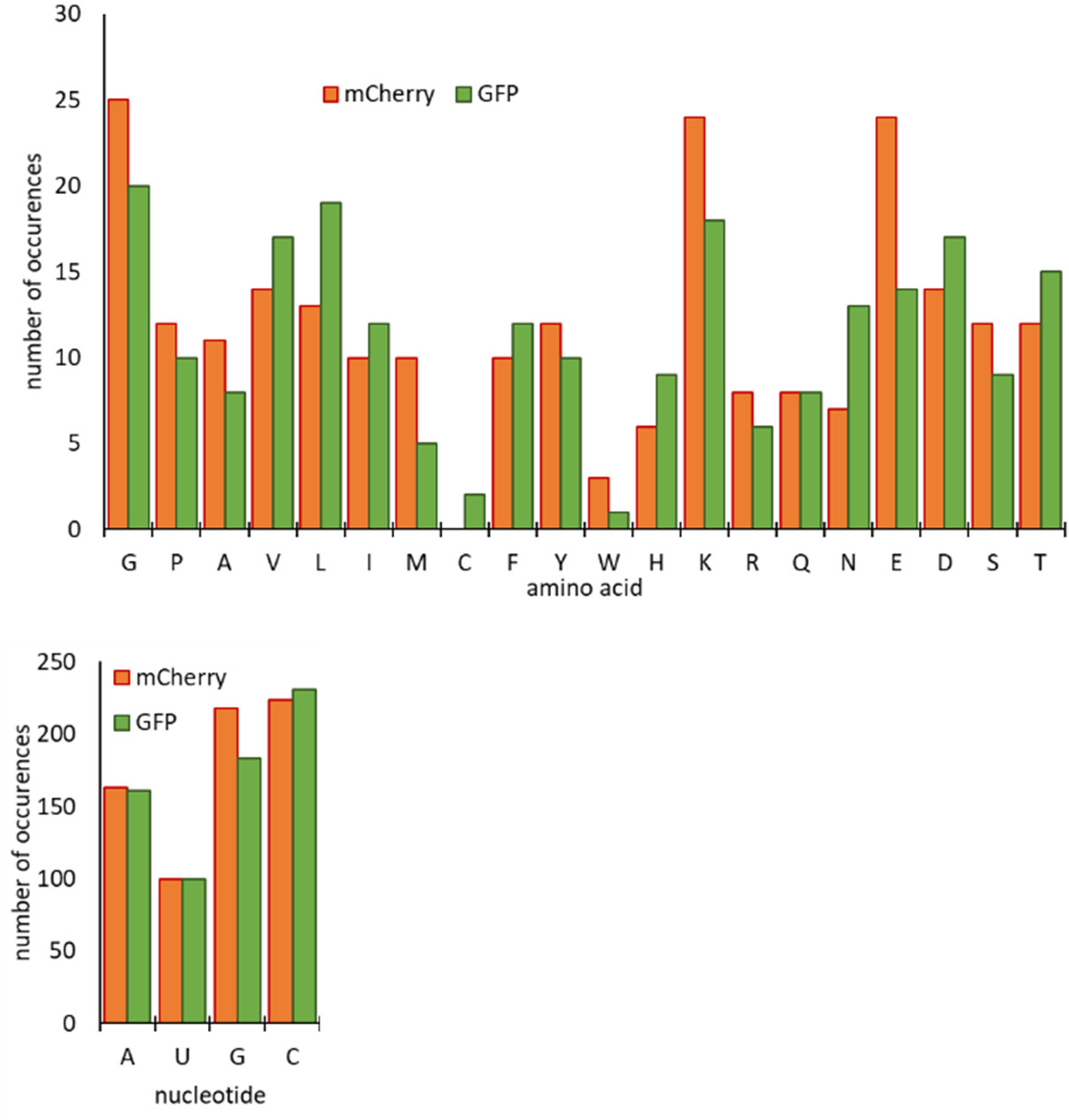
Approximation of resources needed for mCherry and GFP synthesis: the frequency of occurrence of each amino acid in the protein and each nucleotide in the mRNA for both proteins.

**Figure S2.**
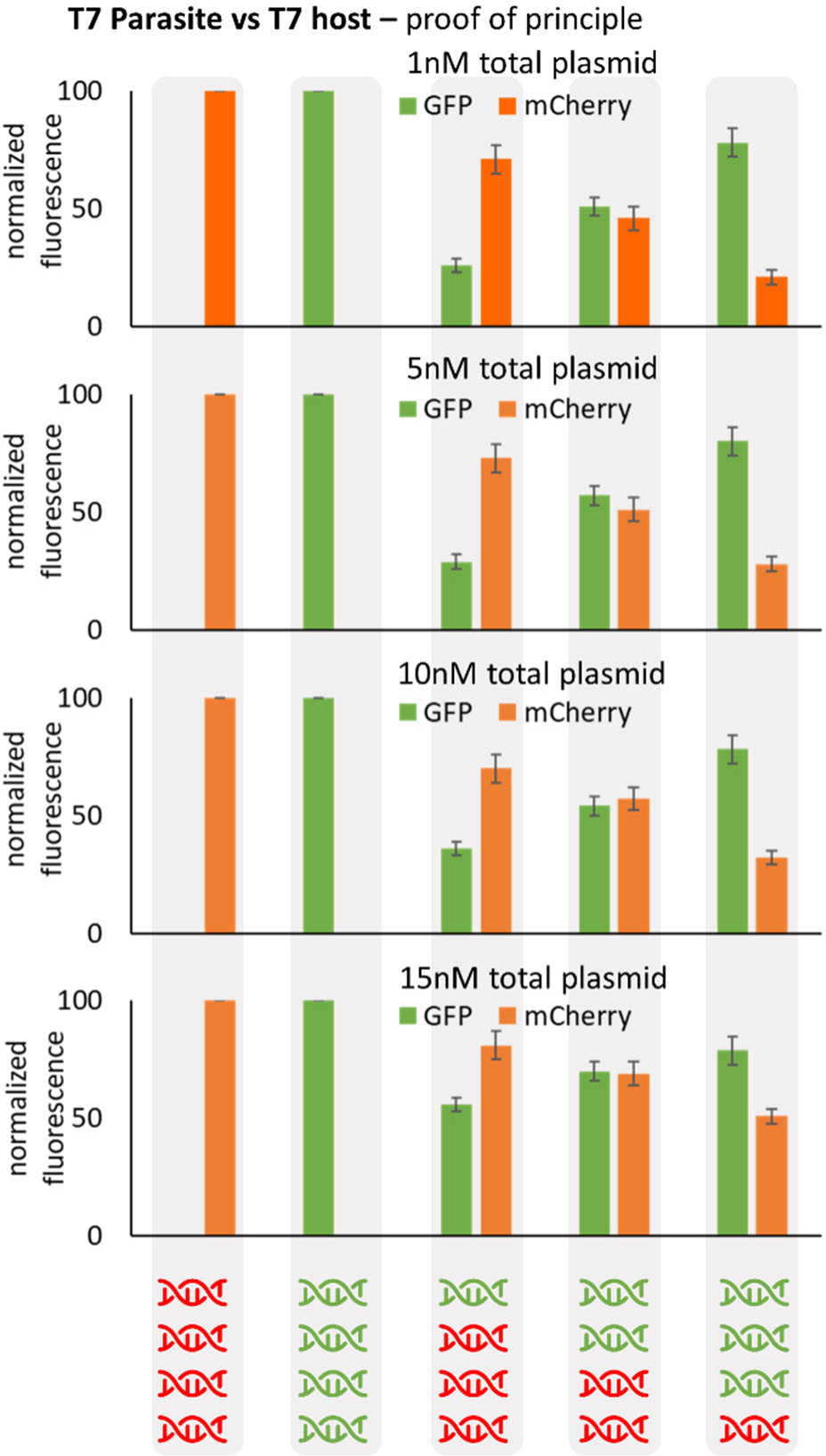
Individual data points for test of expression of plasmids for “parasite” mCherry under T7 promoter and “host” GFP under T7 promoter. Plasmids were mixed at ratios and final plasmid concentrations as indicated on the graph, and expressed in TxTl (without encapsulation). Error bars are SEM n=3.

**Figure S3.**
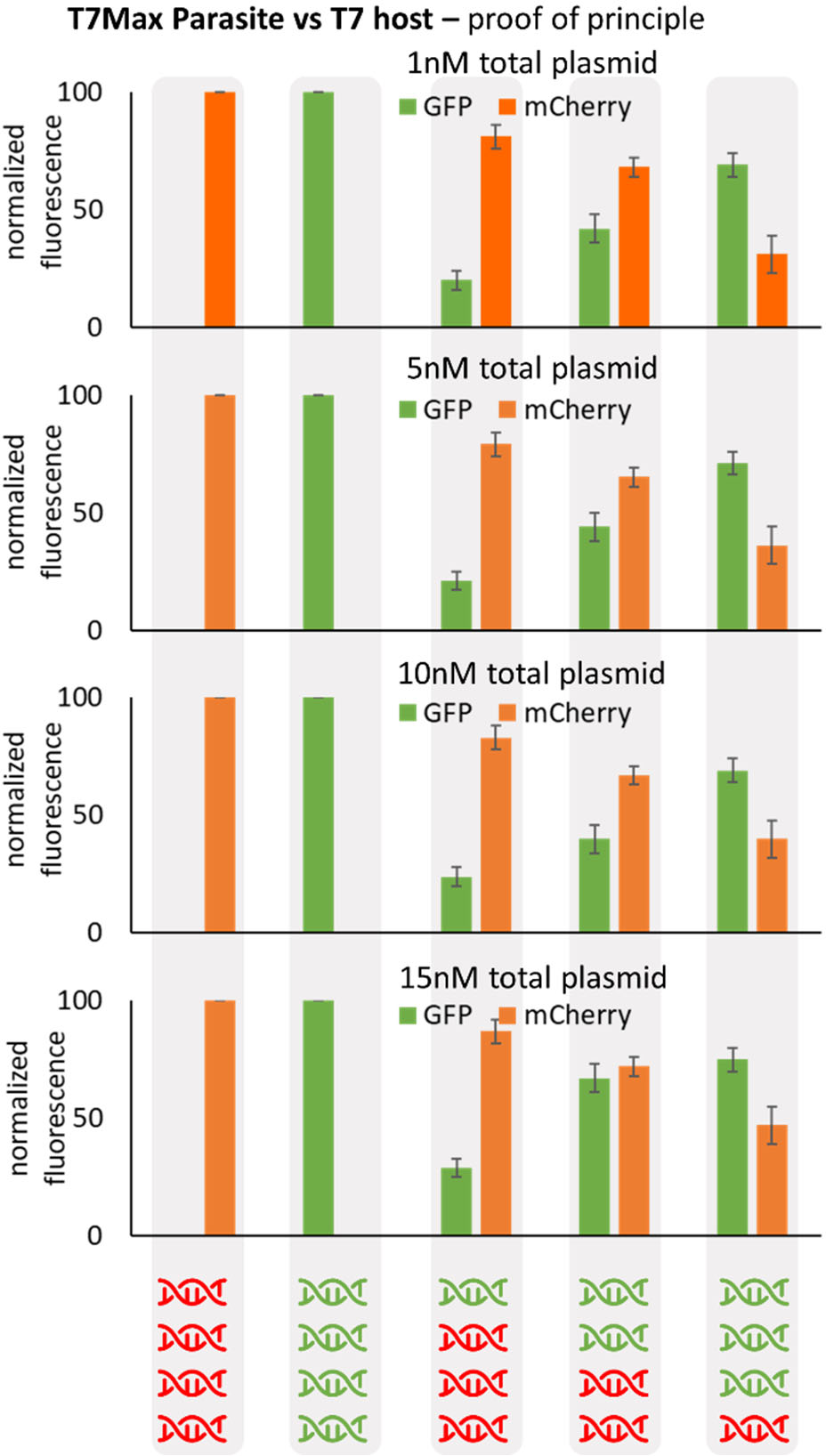
Individual data points for test of expression of plasmids for “parasite” mCherry under T7Max promoter and “host” GFP under T7 promoter. Plasmids were mixed at ratios and final plasmid concentrations as indicated on the graph, and expressed in TxTl (without encapsulation). Error bars are SEM n=3.

**Figure S4.**
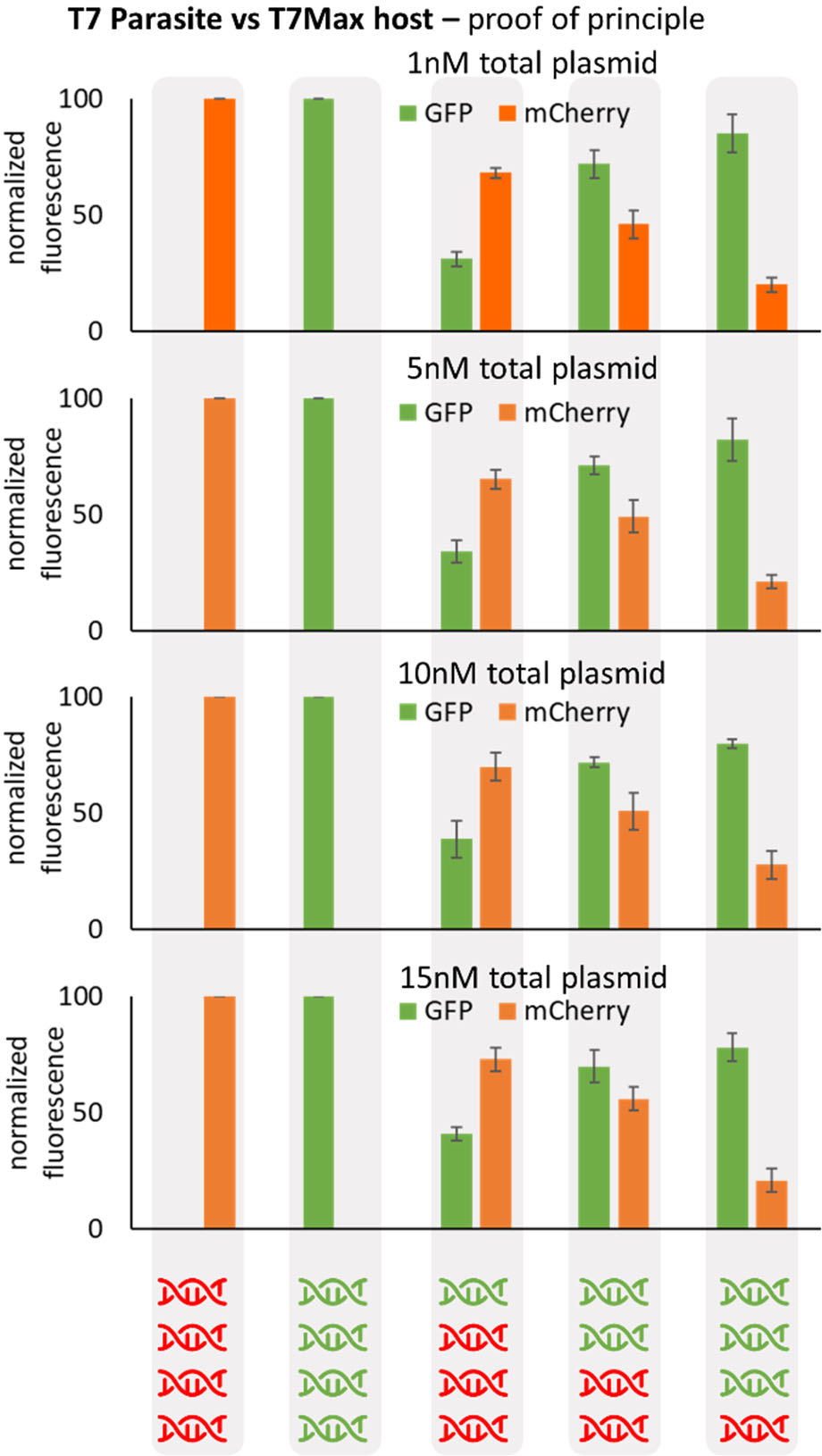
Individual data points for test of expression of plasmids for “parasite” mCherry under T7 promoter and “host” GFP under T7Max promoter. Plasmids were mixed at ratios and final plasmid concentrations as indicated on the graph, and expressed in TxTl (without encapsulation). Error bars are SEM n=3.

**Figure S5.**
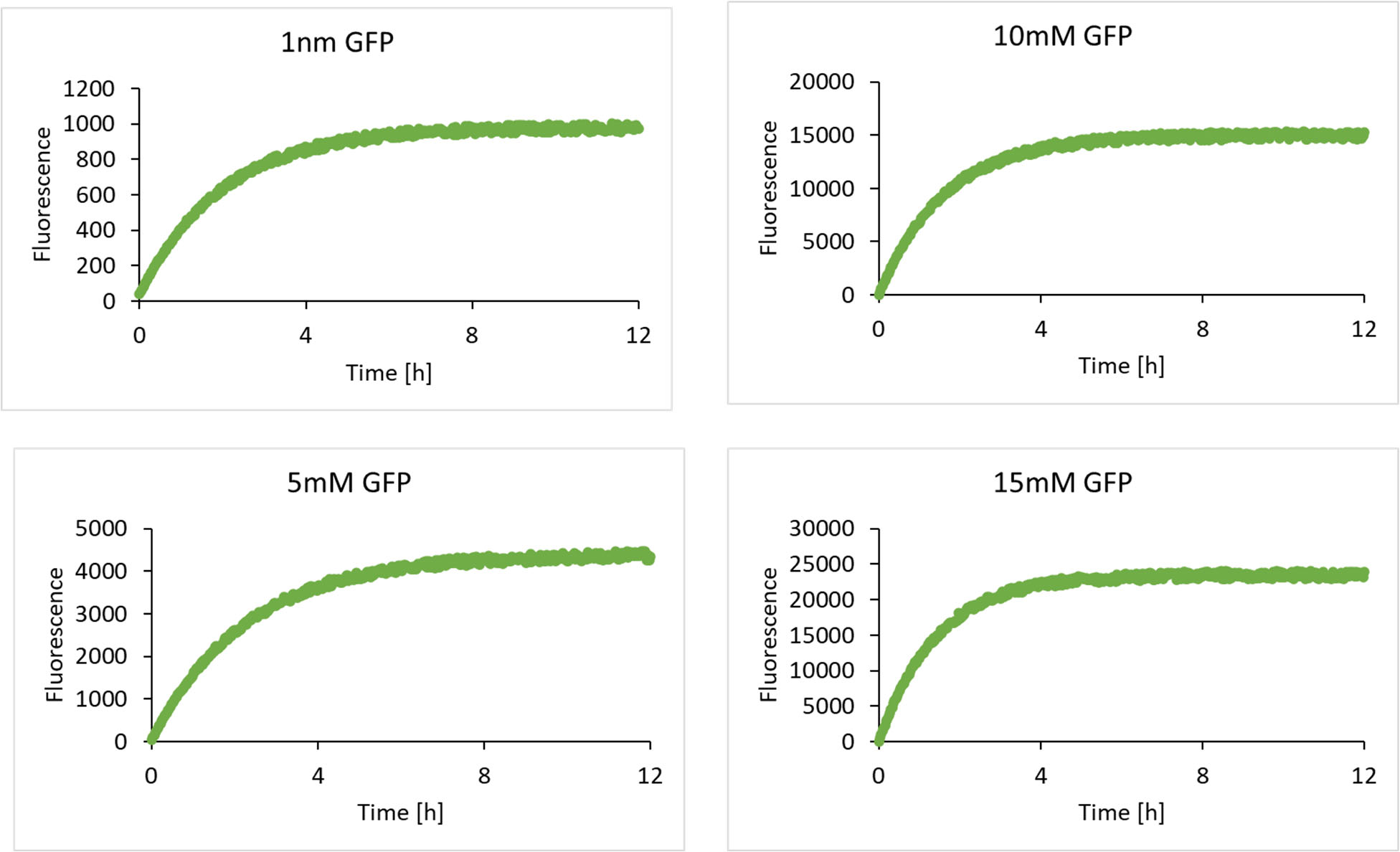
Representative time course of GFP expression for end point data shown on Figure 2b.

**Figure S6.**
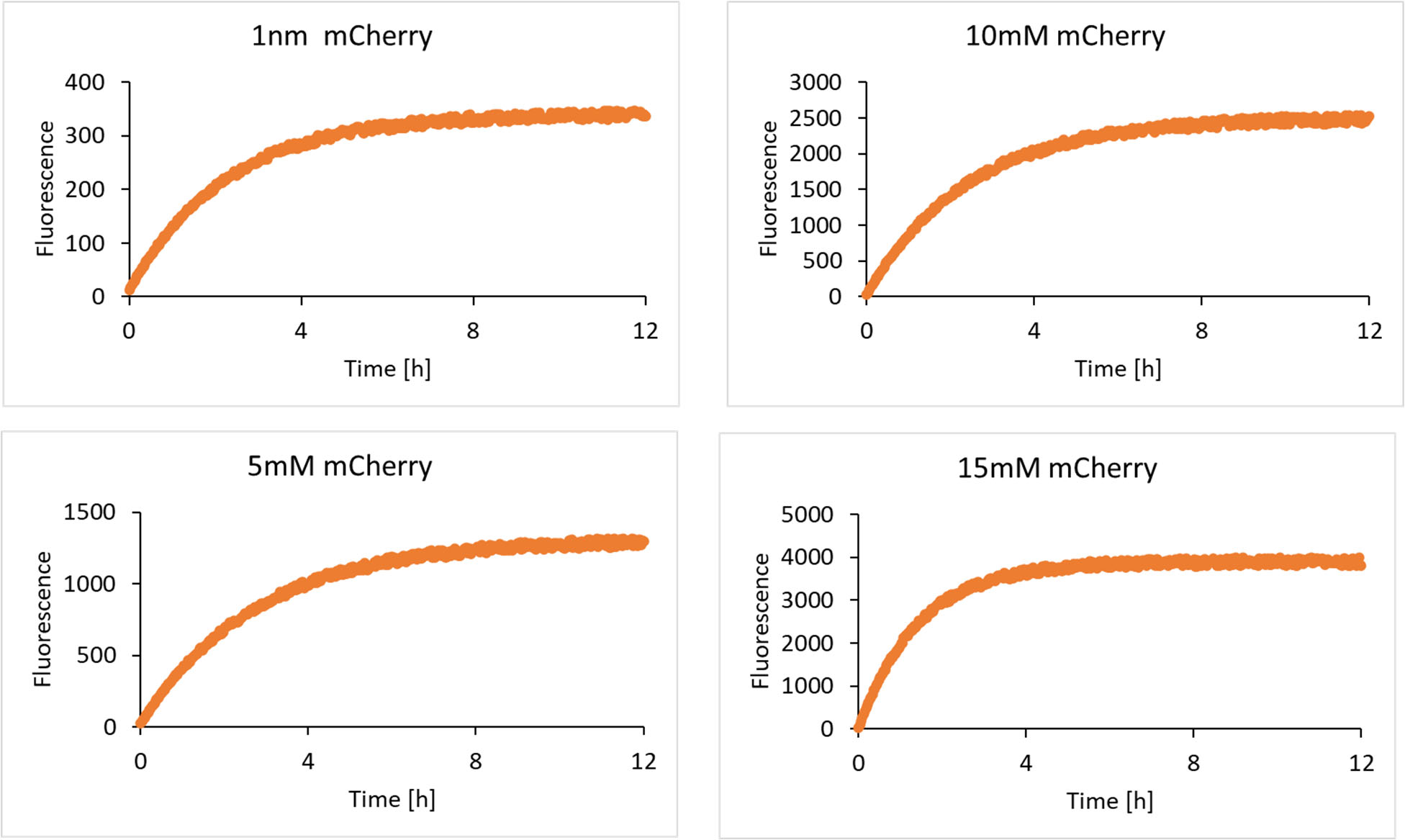
Representative time course of mCherry expression for end point data shown on Figure 2b.

**Figure S7.**
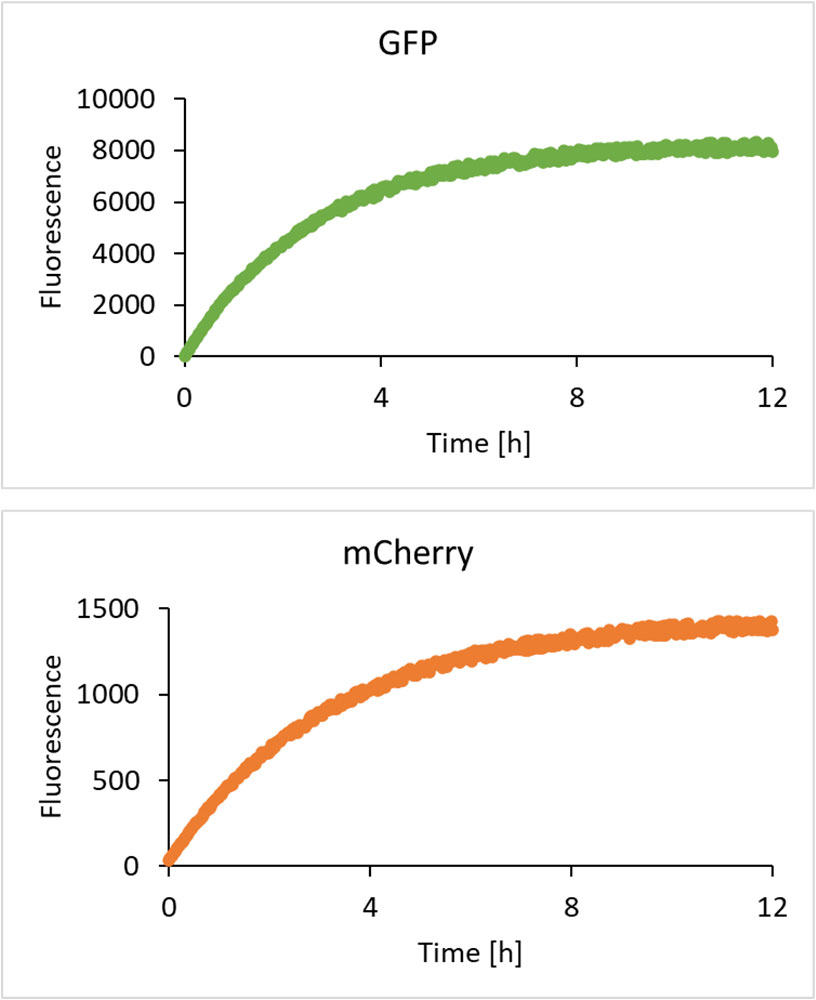
Representative time course of GFP and mCherry expression for end point data shown on Figure 2b, in experiment with 10nM total plasmid, 2G2C (2x GFP and 2x mCherry plasmids).

**Figure S8.**
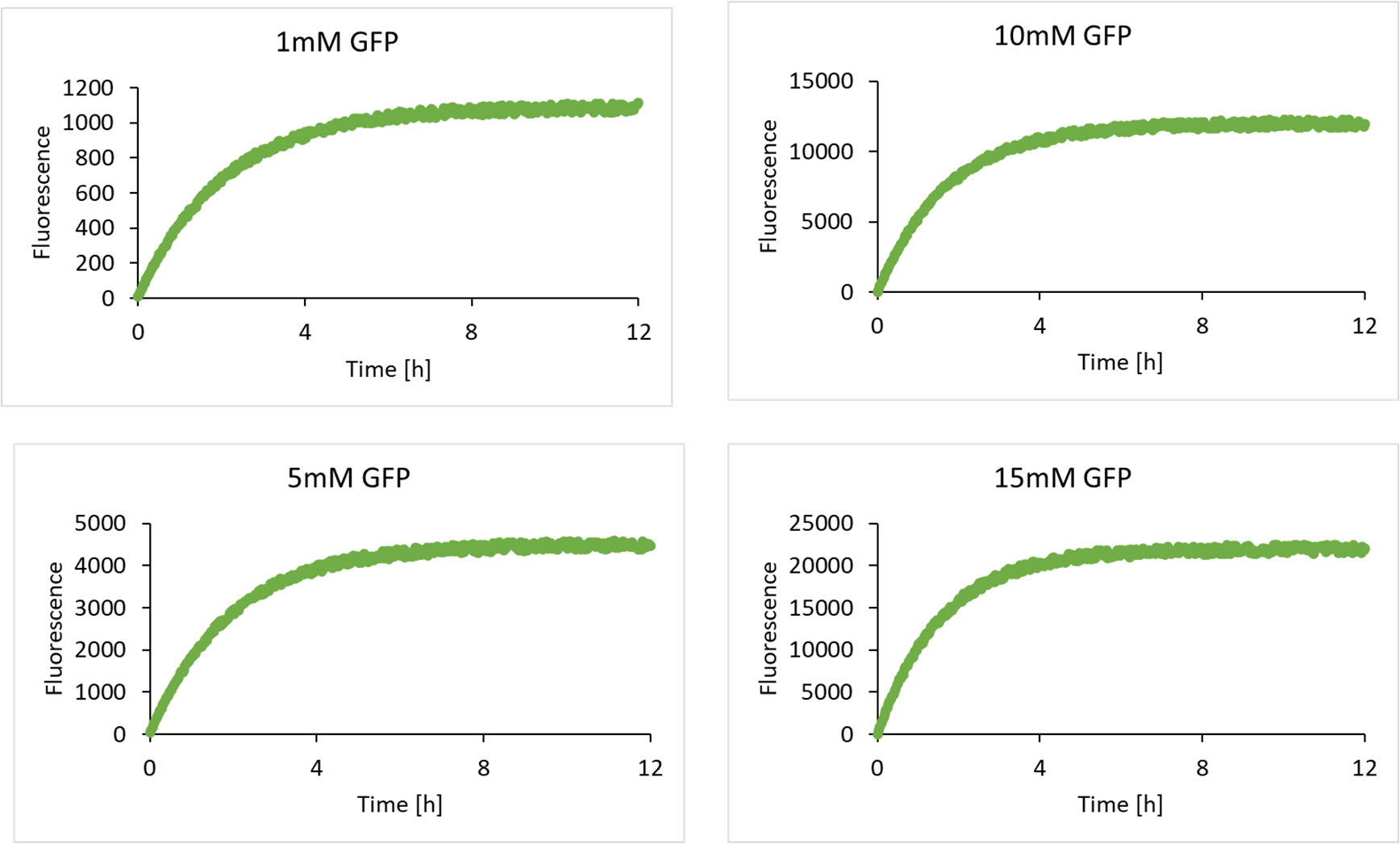
Representative time course of GFP expression for end point data shown on Figure 2c.

**Figure S9.**
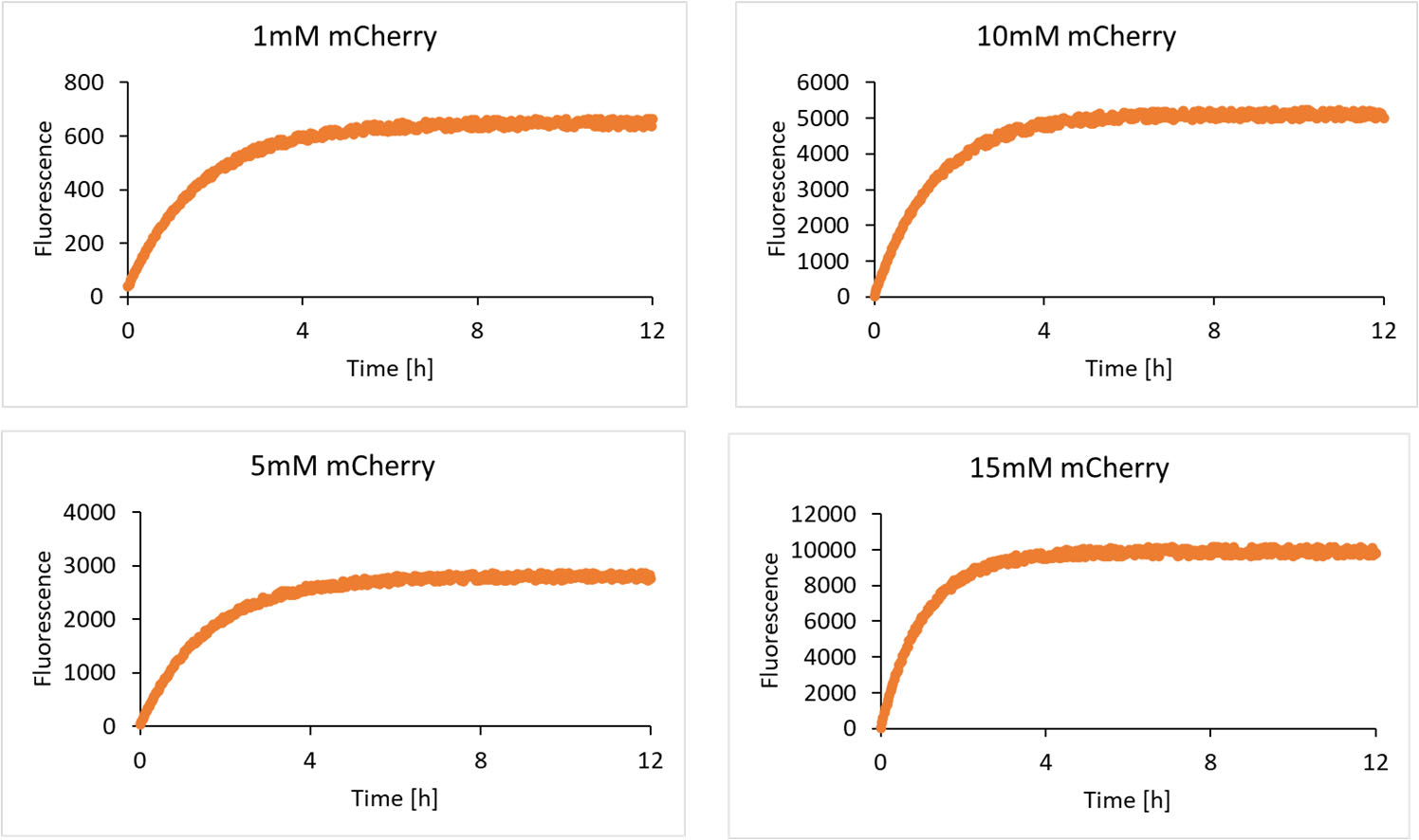
Representative time course of mCherry expression for end point data shown on Figure 2c.

**Figure S10.**
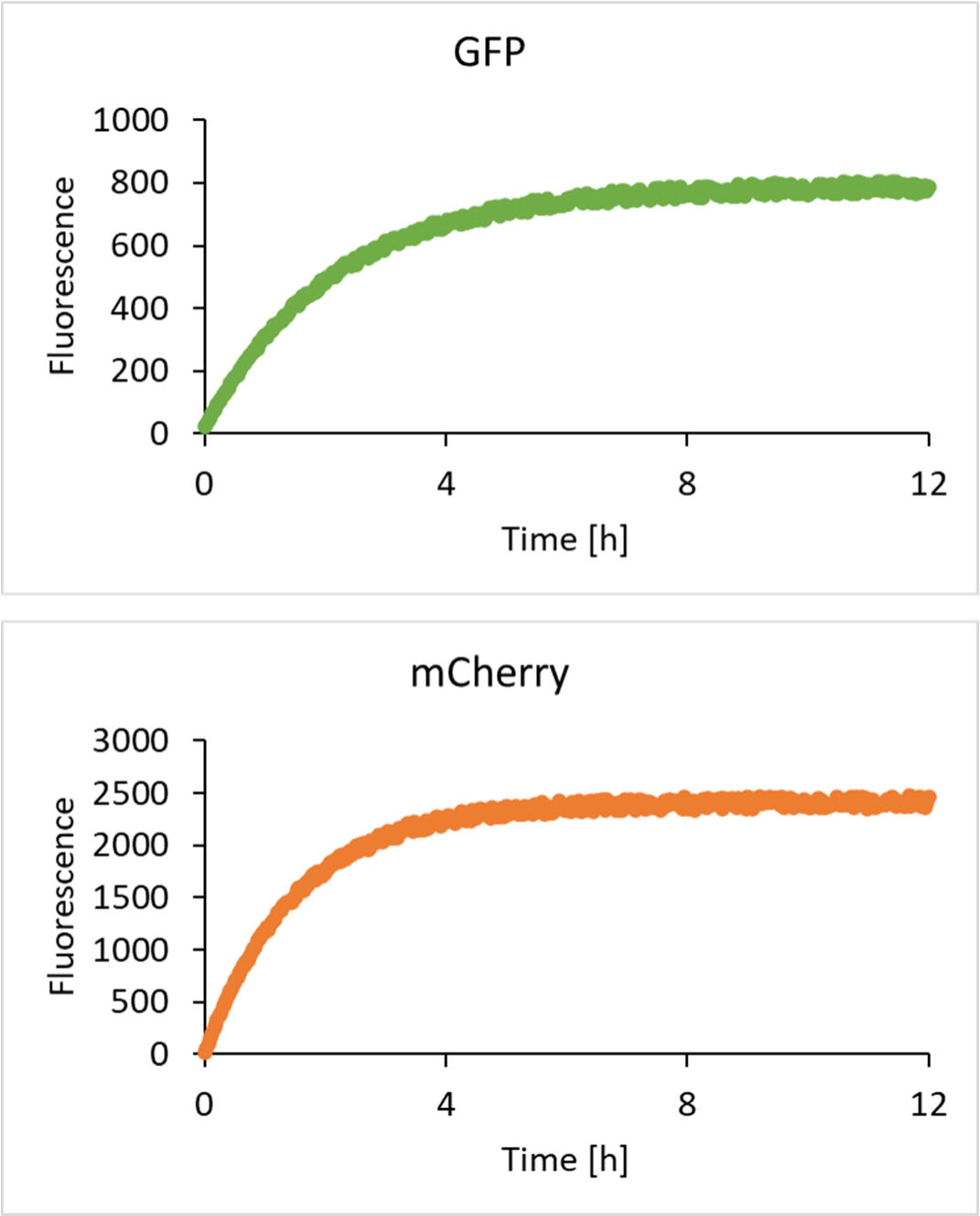
Representative time course of GFP and mCherry expression for end point data shown on Figure 2c, in experiment with 10nM total plasmid, 1G3C (1x GFP and 3x mCherry plasmids).

**Figure S11.**
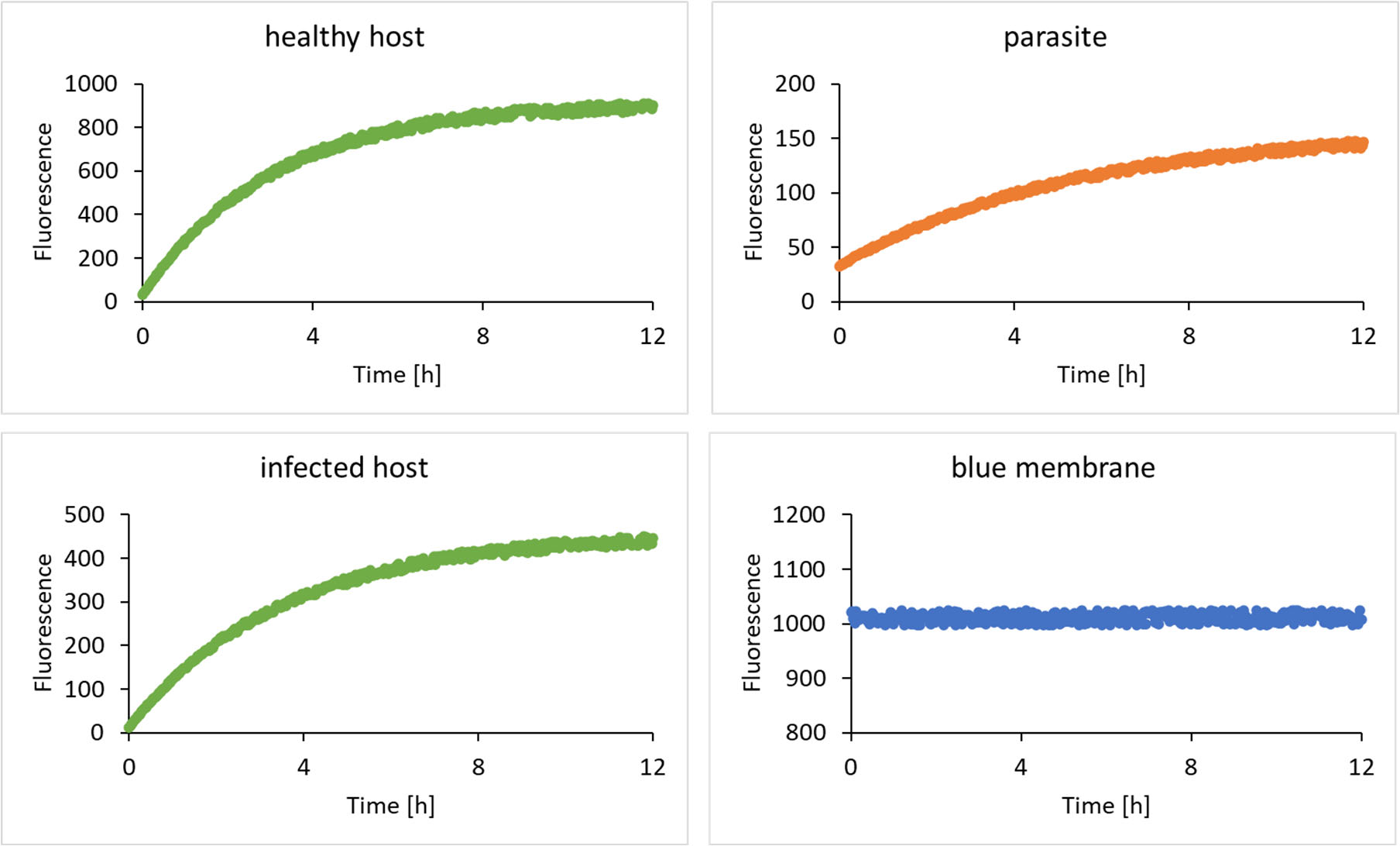
Example time course of GFP and mCherry expression for end point data shown on Figure 2e, in experiment with 1nM total plasmid, equal amount of host (GFP) and parasite (mCherry). Blue trace is membrane dye Marina Blue DHPE.

**Figure S12.**
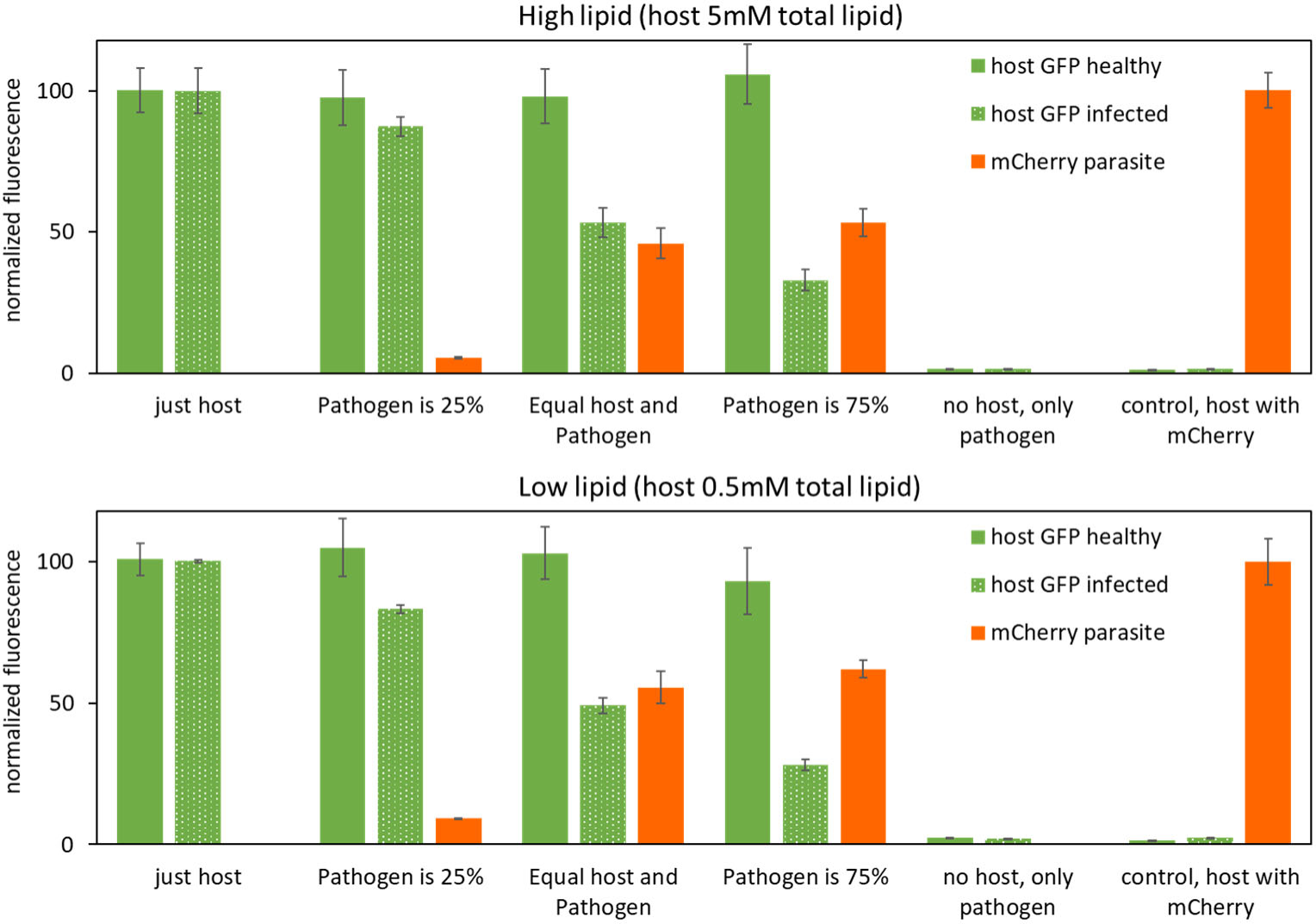
Parasite infection according to scheme shown on figure 2c, with higher and lower total lipid concentration (preserving the host and parasite lipid ratios). Those experiments were intended to establish that no lipid concentration change artifacts are possibly responsible for the observed results. The experiments were started with host concentration of 0.5mM and 5mM total lipid, respectively, for low and high lipid concentration samples.

**Figure S13.**
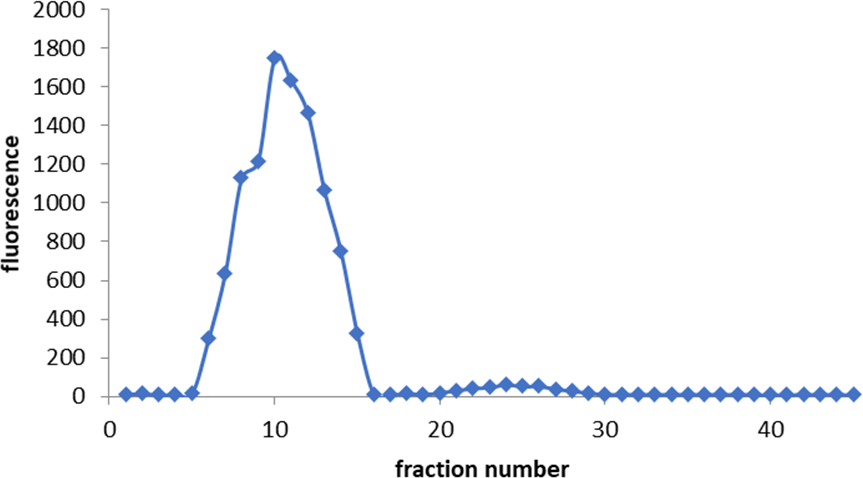
Liposome samples purified on Sepharose size exclusion column after “infection” experiments with liposome fusion. GFP fluorescence is measured in green channel. In this experiment, parasite is 75% total population.

**Figure S14.**
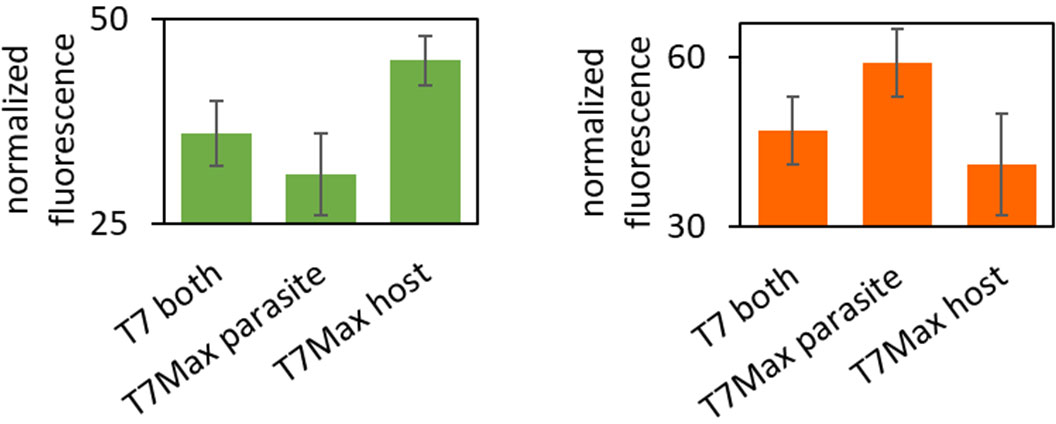
Side by side comparison of host (GFP, green) and parasite (mCherry, red) for infected populations with parasite being 75% total population – the values signified with the star on figure 2. Error bars are SEM n=3.

**Figure S15.**
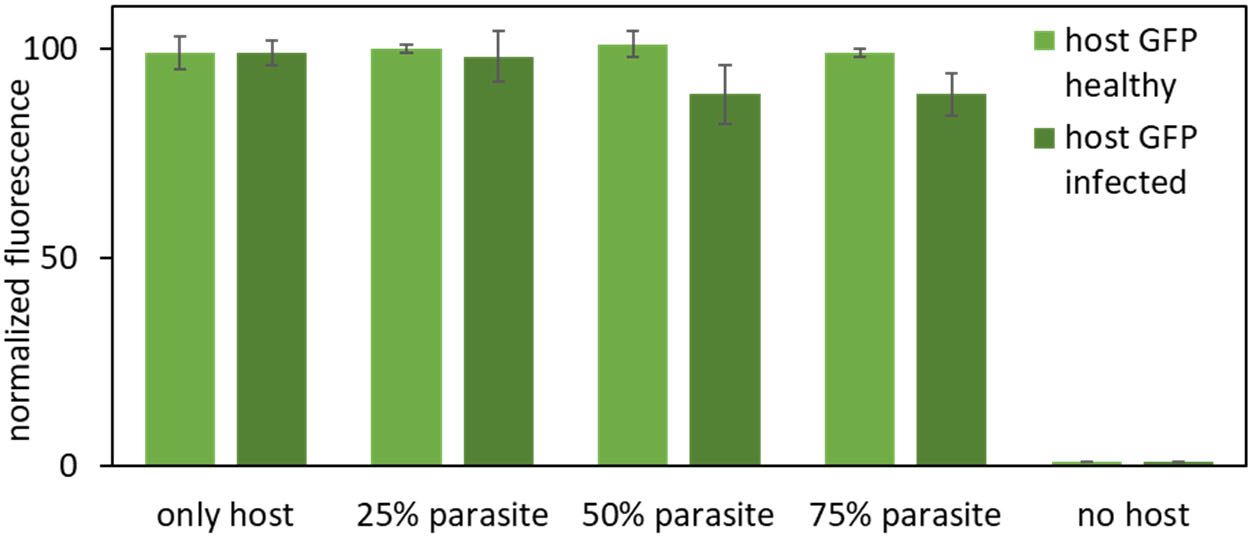
Infection with “parasite” cells that lack plasmid, under the same conditions are results shown on figure 2h.

**Figure S16.**
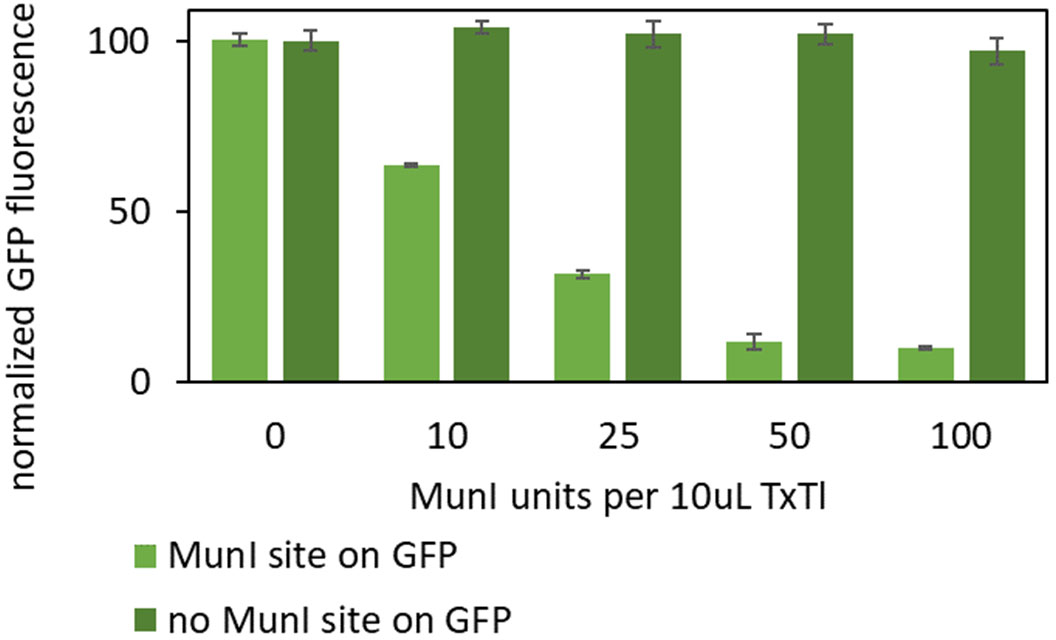
Proof of principle for MunI infection experiments: 5mM GFP plasmid in TxTl with MunI enzyme added outside. The enzyme used, Thermo Scientific MunI (Catalog number: ER0751) was supplied at concentration of 10 U/µL. According to the manufacturer: “One unit is defined as the amount of MunI required to digest 1 μg of lambda DNA in 1 hour at 37°C in 50 μL of recommended reaction buffer.”. The protein concentration for the enzyme was not available. Individual time courses for the experiments are on **Figure S17**.

**Figure S17.**
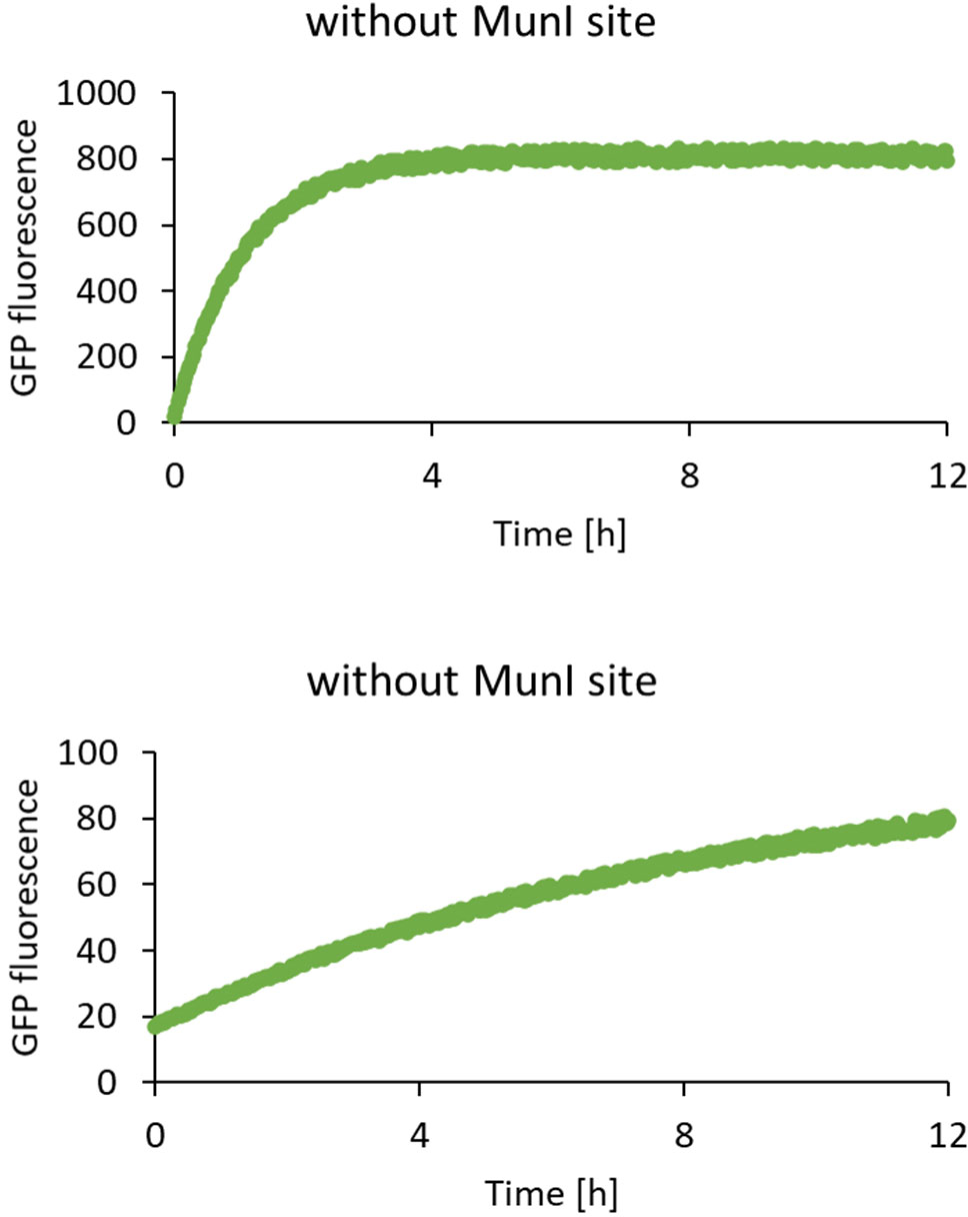
Individual protein expression time course data for samples with 50U of MunI purified enzyme, using GFP plasmid with and without MunI restriction site.

**Figure S18.**
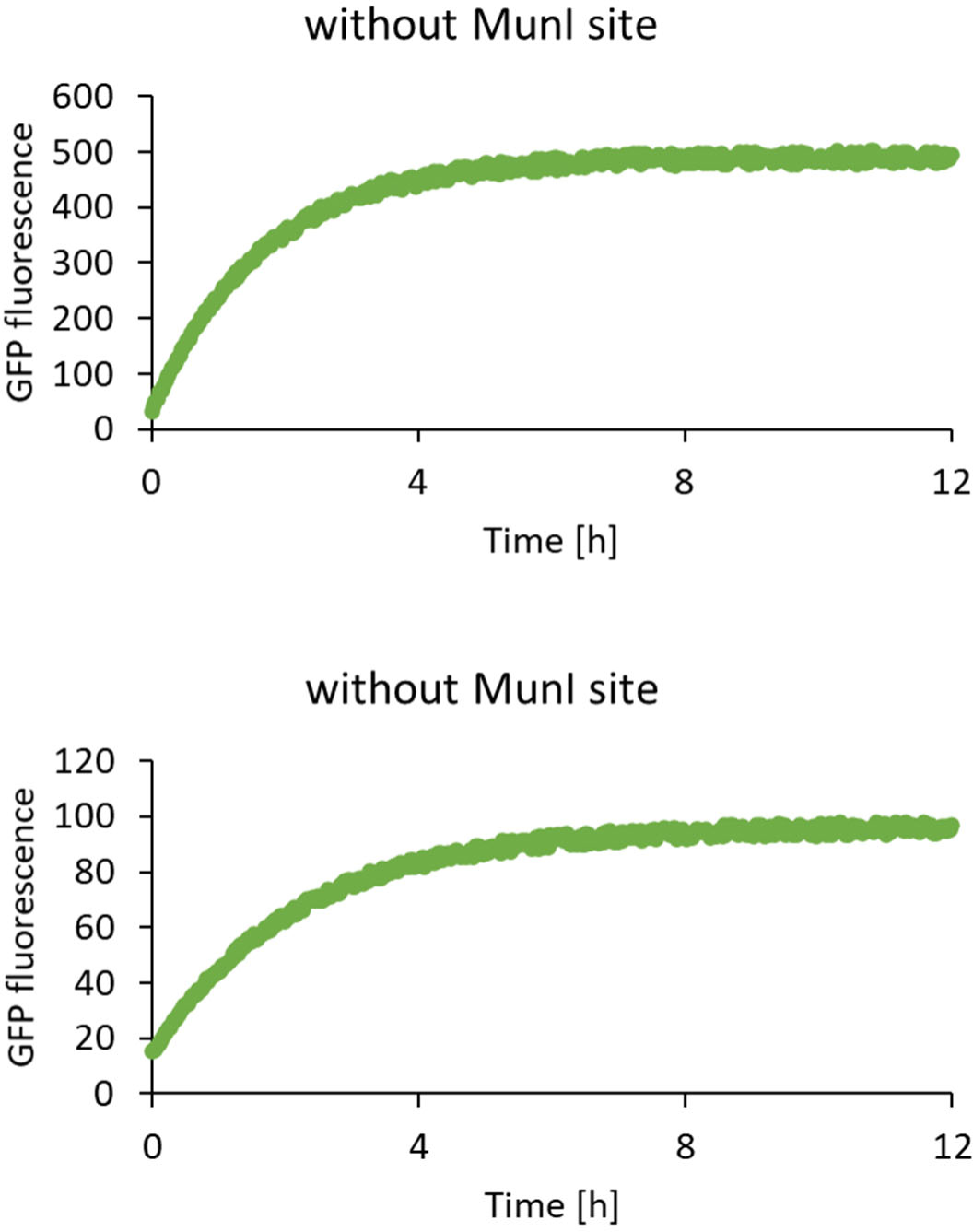
Individual protein expression time course data for samples with 5nM of MunI plasmid, using GFP plasmid with and without MunI restriction site.

**Figure S19.**
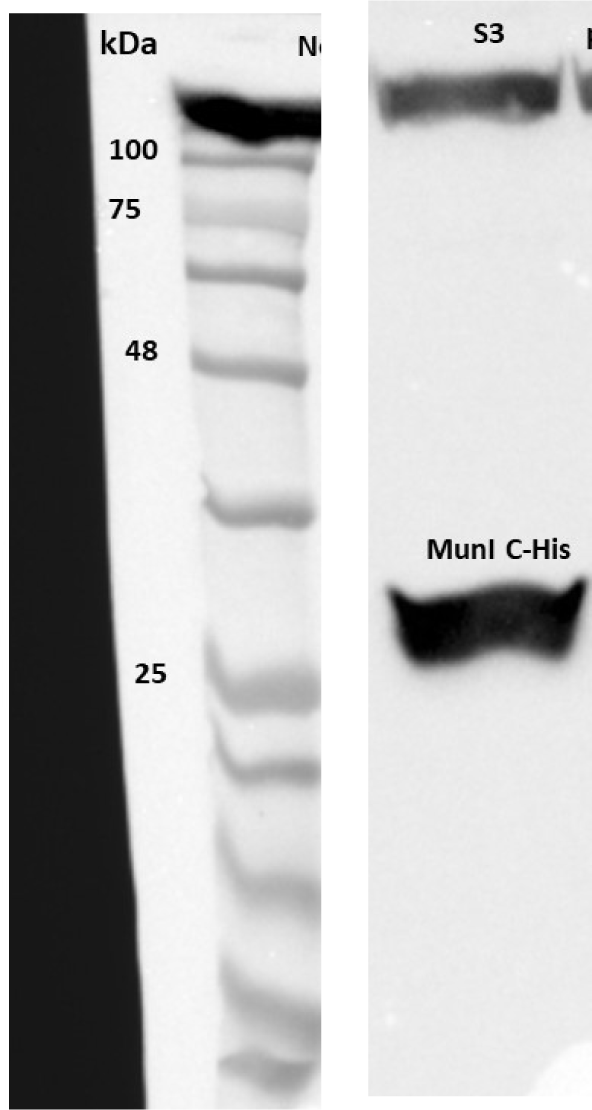
Example Western Blot of MunI restriction enzyme with 6xHis tag, 5nM MunI plasmid. The ladder is a BLUEstain 2 Protein Ladder. The digest activity of MunI enzyme expressed in TxTl is shown on Figure S33.

**Figure S20.**
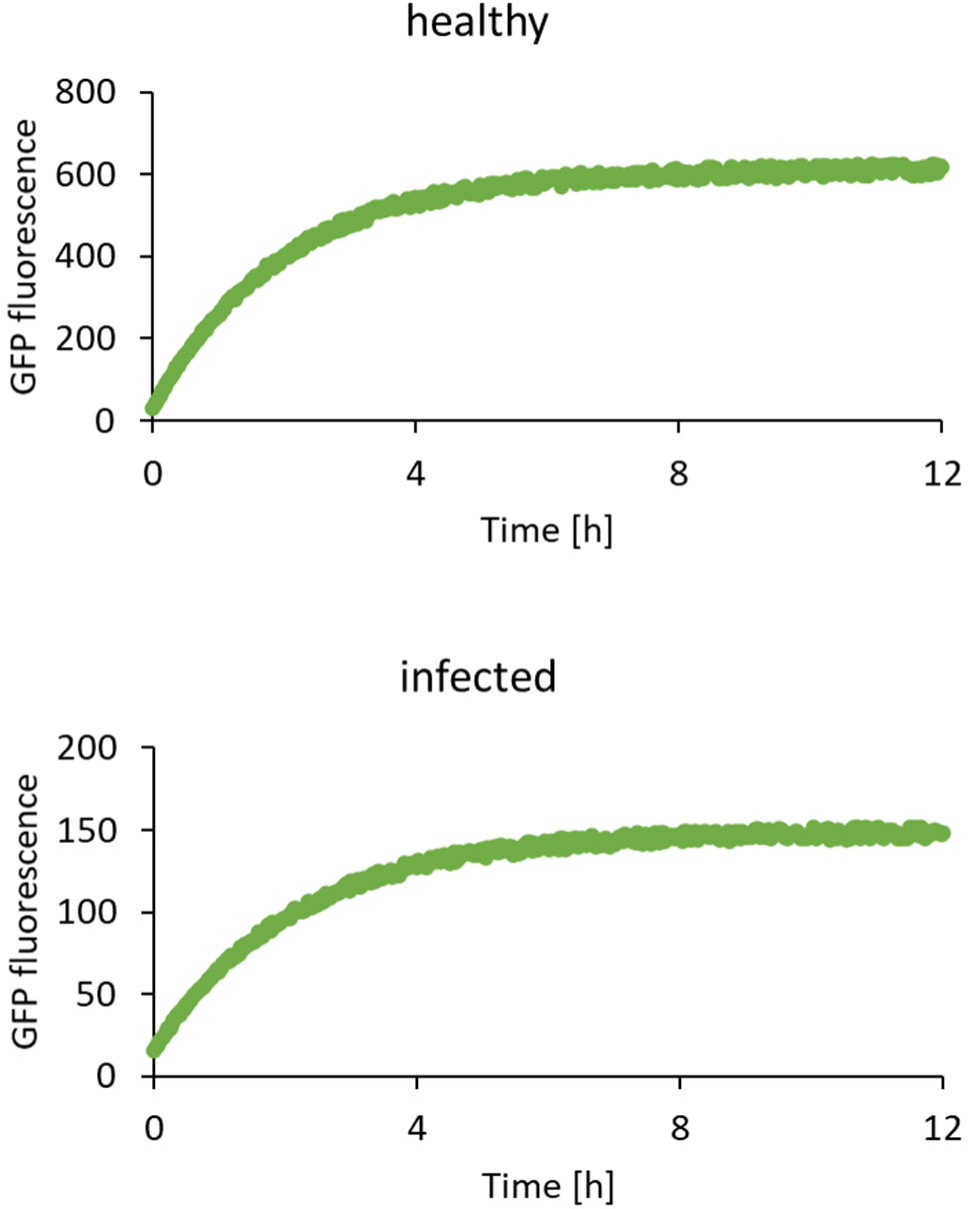
Individual protein expression time course data for samples with Pathogen being 75% of the population, for “infected” and “healthy” GFP expression.

**Figure S21.**
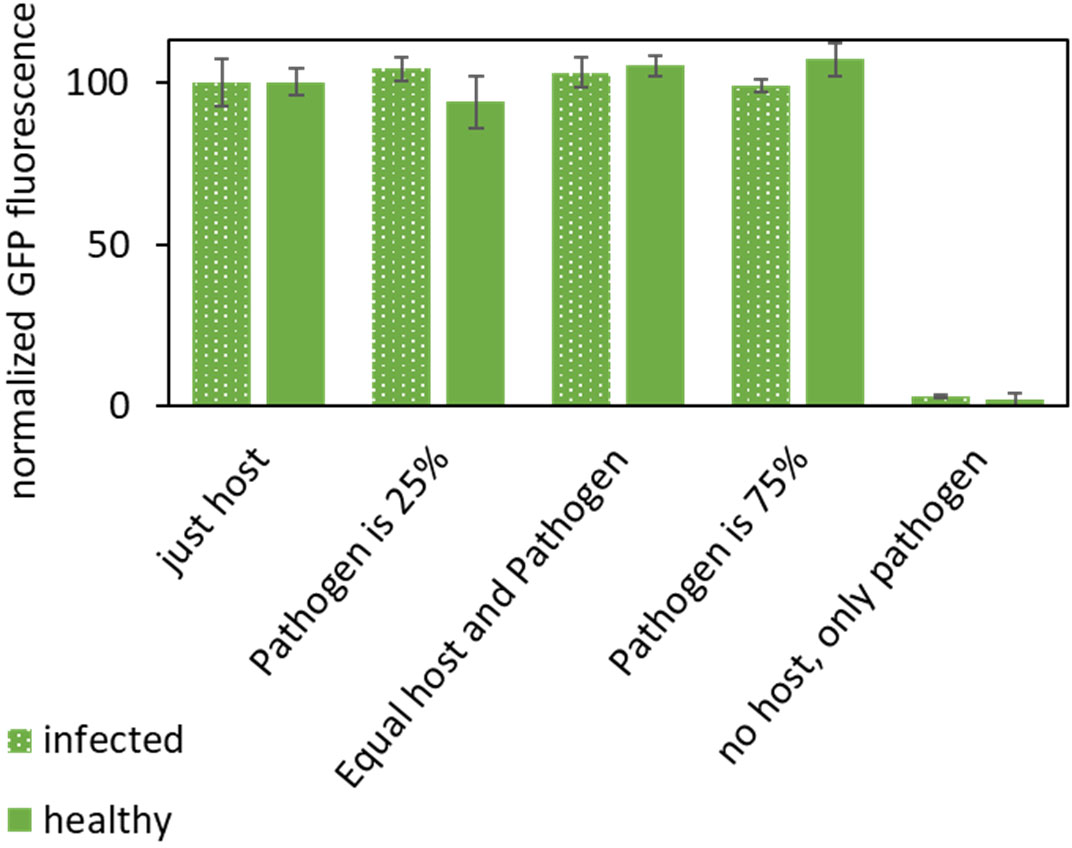
Infection with incompatible fusion tags on host and parasite. The experiments were performed the same way as experiments shown on Figure 3d, except the sequences of the DNA fusion tags on the surface of the liposomes were not compatible fusion partners.

**Figure S22.**
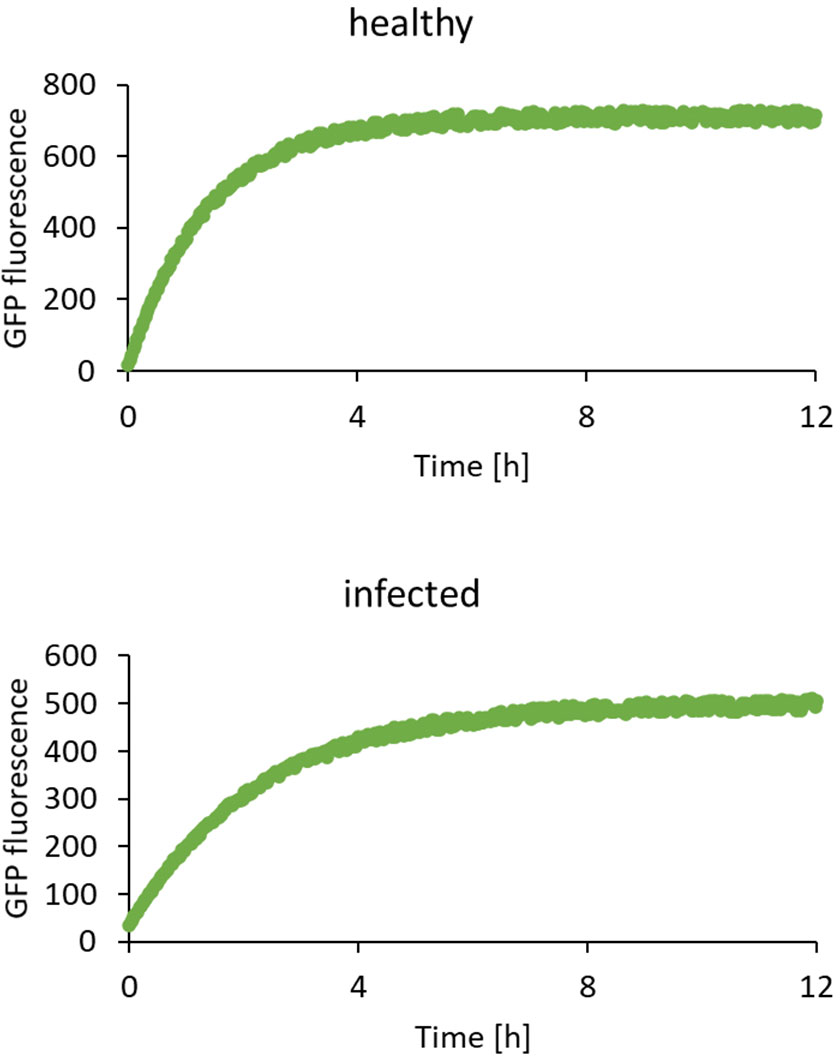
Individual protein expression time course data for samples with Pathogen being 75% of the population, for “infected” and “healthy” GFP expression, with host “immune” to the pathogen (no MunI site in the GFP plasmid).

**Figure S23.**
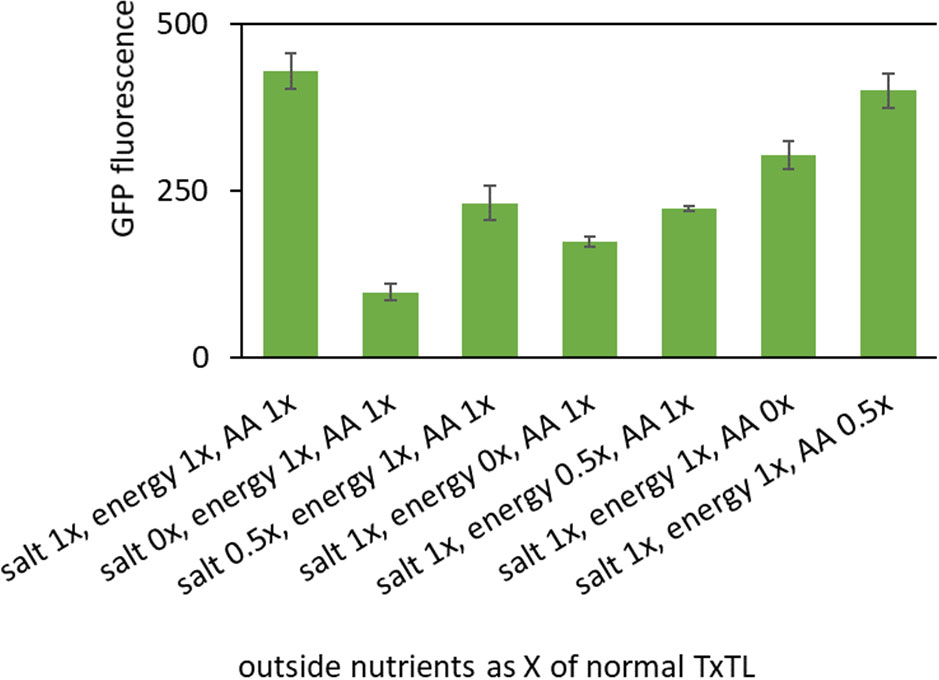
Nutrient concentration in aHL rescue experiment. All samples had 10nM aHL plasmid and 10nM GFP plasmid.

**Figure S24.**
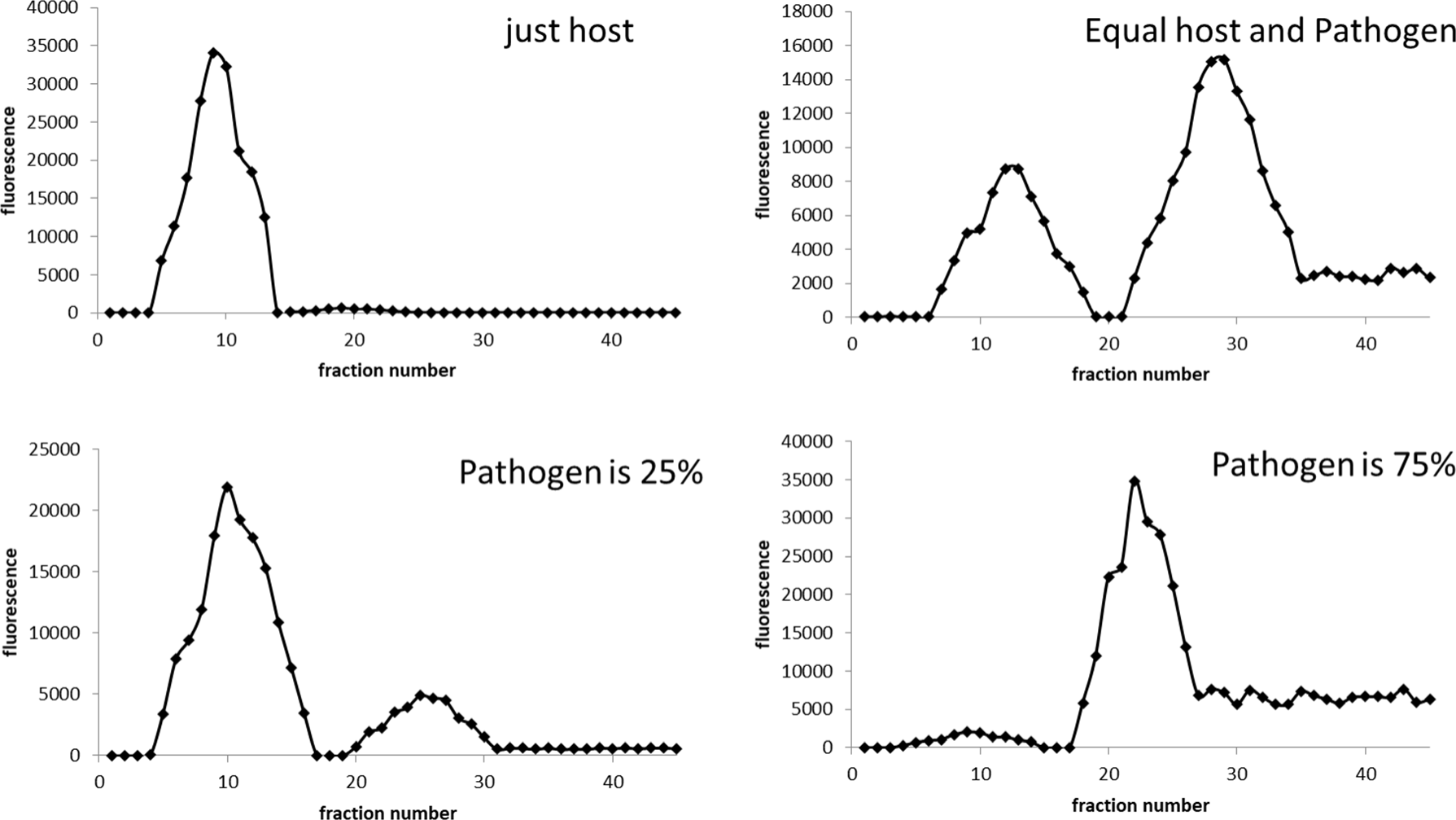
Leakage of small molecules from infection of aHL parasite. Samples were prepared identical as in experiments demonstrated on figure 4c, with two changes: host had mCherry fluorescent protein instead of GFP, and 0.5mM calcein was encapsulated inside host cells. After reaction, liposome samples were purified on Sepharose size exclusion purification column as previously described.^30^ The small molecule presence inside and outside of liposomes was detected in the green channel (calcein). All experiments on this figure are for the “infected” scenario: host expresses aHL.

**Figure S25.**
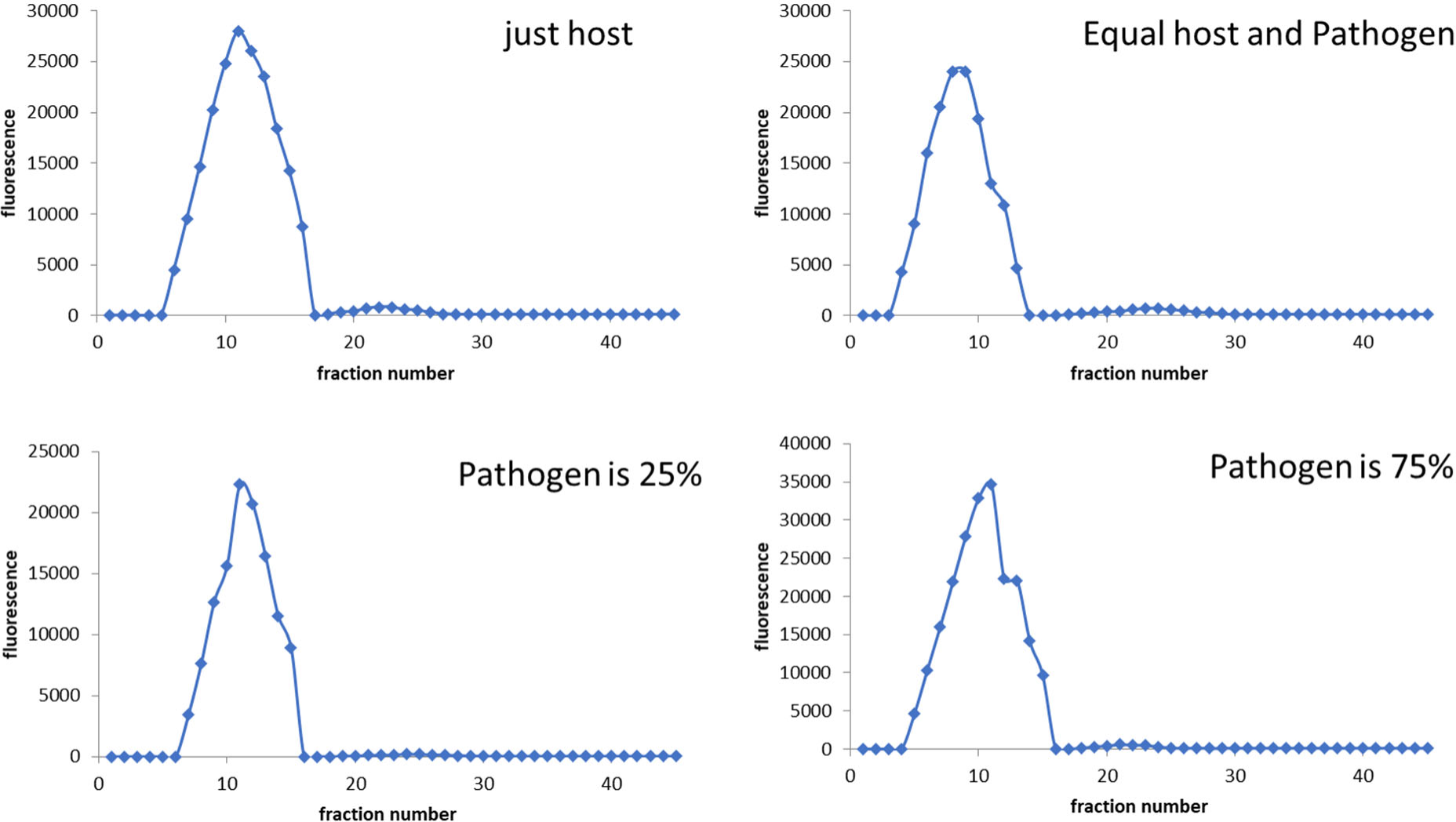
Leakage of small molecules from infection of aHL parasite. Samples were prepared identical as in experiments demonstrated on figure 4c, with two changes: host had mCherry fluorescent protein instead of GFP, and 0.5mM calcein was encapsulated inside host cells. After reaction, liposome samples were purified on Sepharose size exclusion purification column as previously described.^30^ The small molecule presence inside and outside of liposomes was detected in the green channel (calcein). All experiments on this figure are for the “healthy” scenario: host cannot expresses aHL.

**Figure S26.**
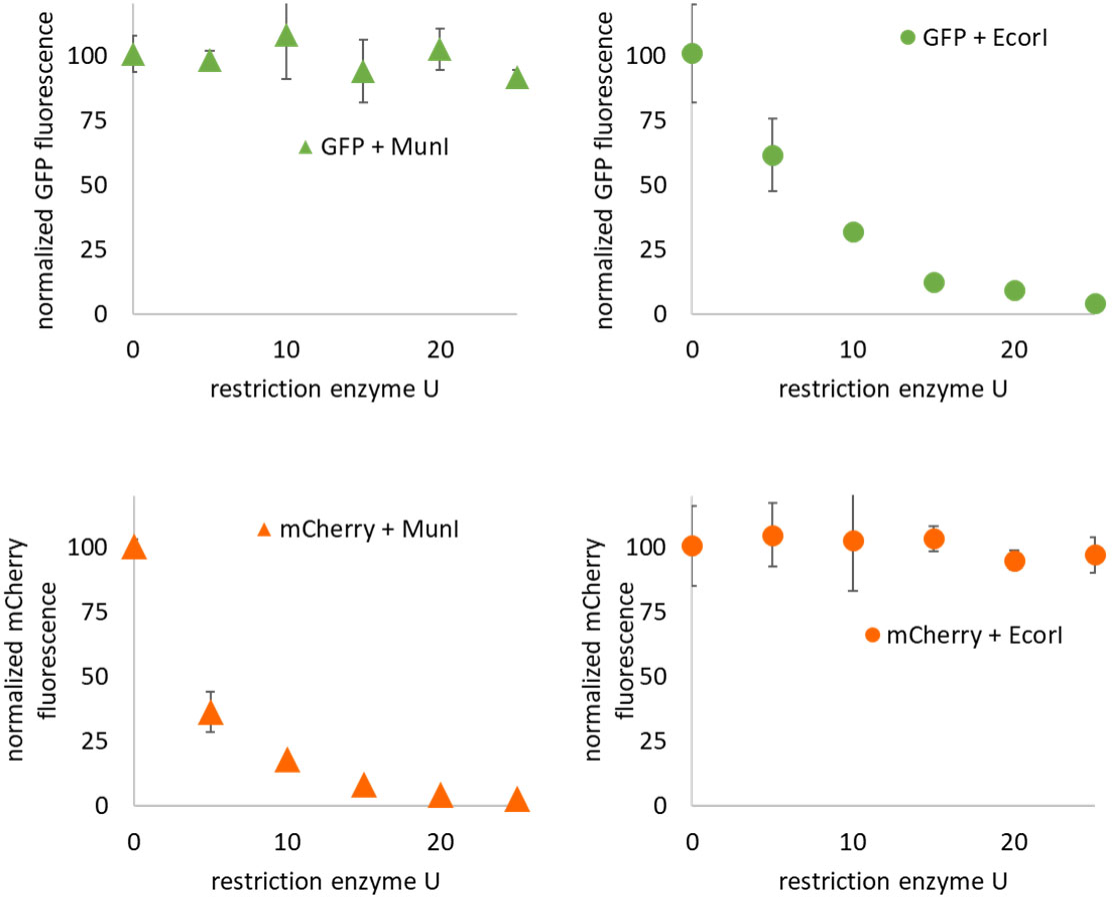
Susceptibility of TxTl plasmid to restriction enzyme digestion, as control for metabolic burden of parasite resistance experiments presented on figure 5b**-e**. The enzymes used, Thermo Scientific MunI (Catalog number: ER0751) and EcoRI (Catalog number: ER0271), were supplied at concentration of 10 U/µL. According to the manufacturer: “One unit is defined as the amount of enzyme (*same for EcorI and MunI*) required to digest 1 μg of lambda DNA in 1 hour at 37°C in 50 μL of recommended reaction buffer.”.

**Figure S27.**
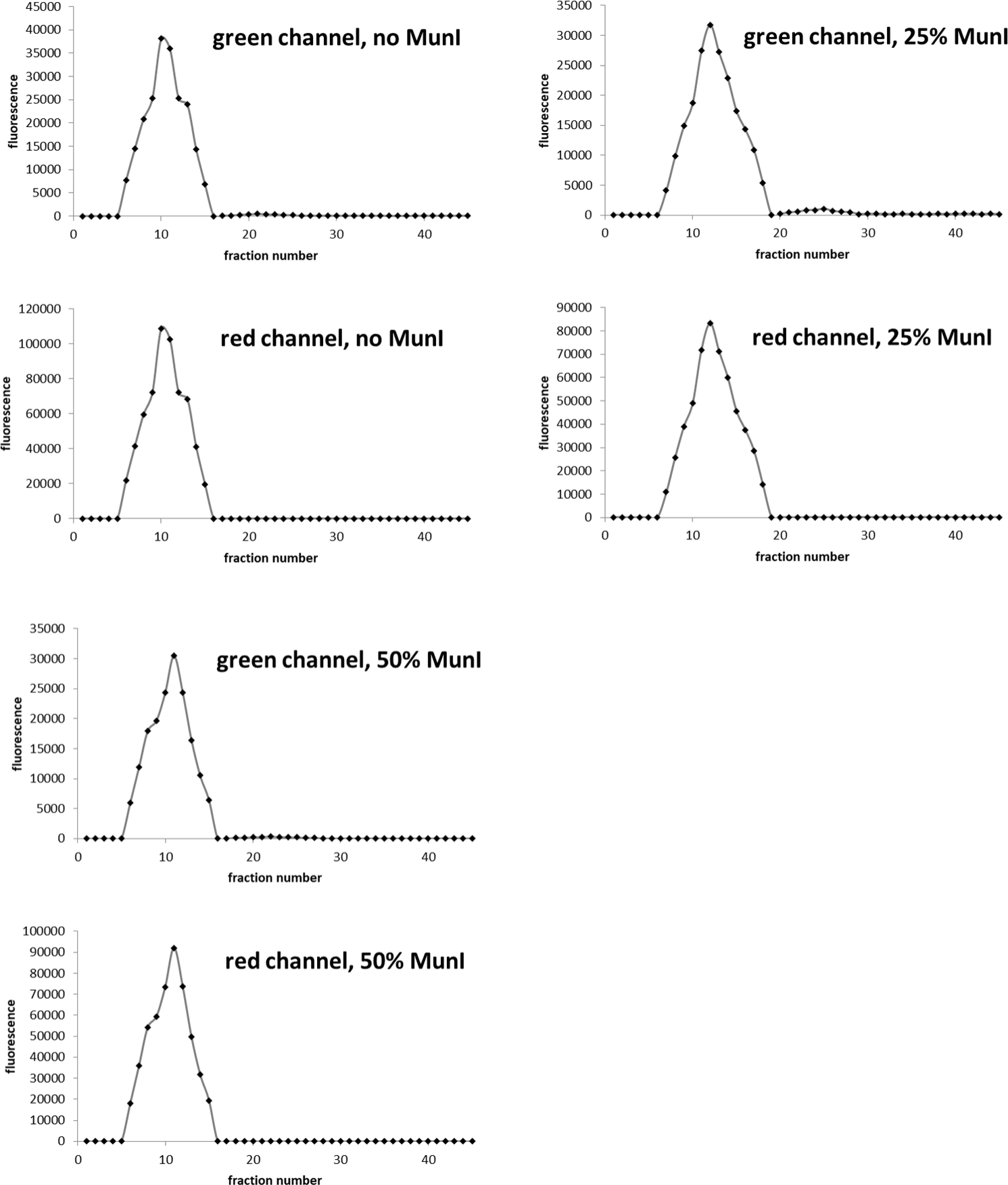
Stability of liposomes expressing increasing amount of parasite (MunI) with GFP. Liposome samples purified on Sepharose size exclusion column after 12h incubation with varying ratio of GFP and MunI plasmids. To increase sensitivity of leakage experiments, liposomes were prepared with calcein, a small molecule dye that will leak out of destabilized membranes easier than large protein like GFP. The green channel shows signal from both GFP and calcein. The red channel indicates position of the liposomes by the red fluorescent membrane dye rhodamine (introduced to the membrane at 0.1mol% as Lissamine™ Rhodamine B 1,2-Dihexadecanoyl-sn-Glycero-3-Phosphoethanolamine, Triethylammonium Salt).

**Figure S28.**
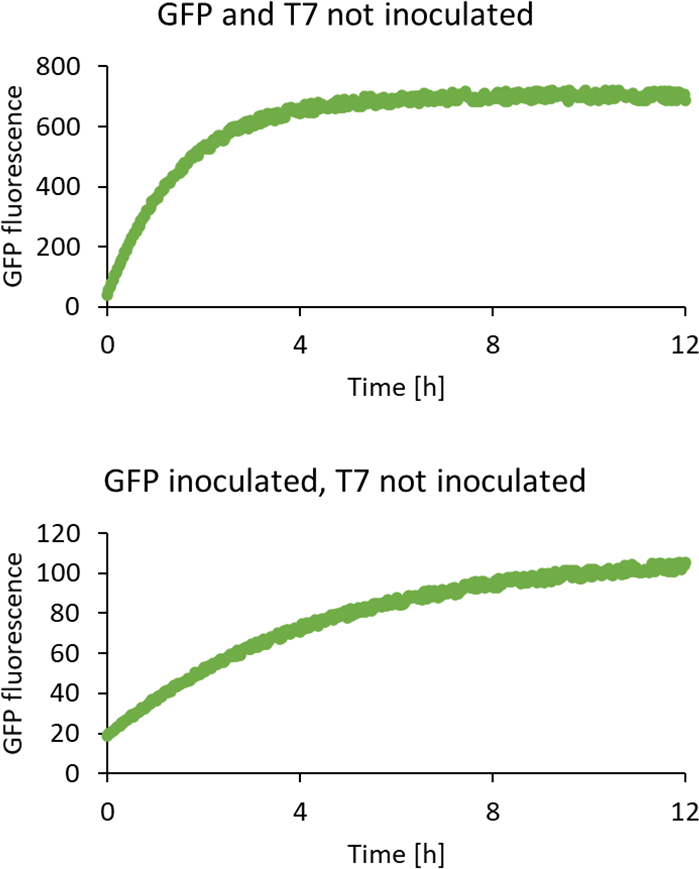
Representative time course for GFP expression in “inoculation” test. Two synthetic cell populations with compatible fusion tags, one containing GFP gene under T7 promoter and the other one containing T7 RNA polymerase, were mixed in equal proportions. One or both of the mixing partners were “inoculated” before mixing.

**Figure S29.**
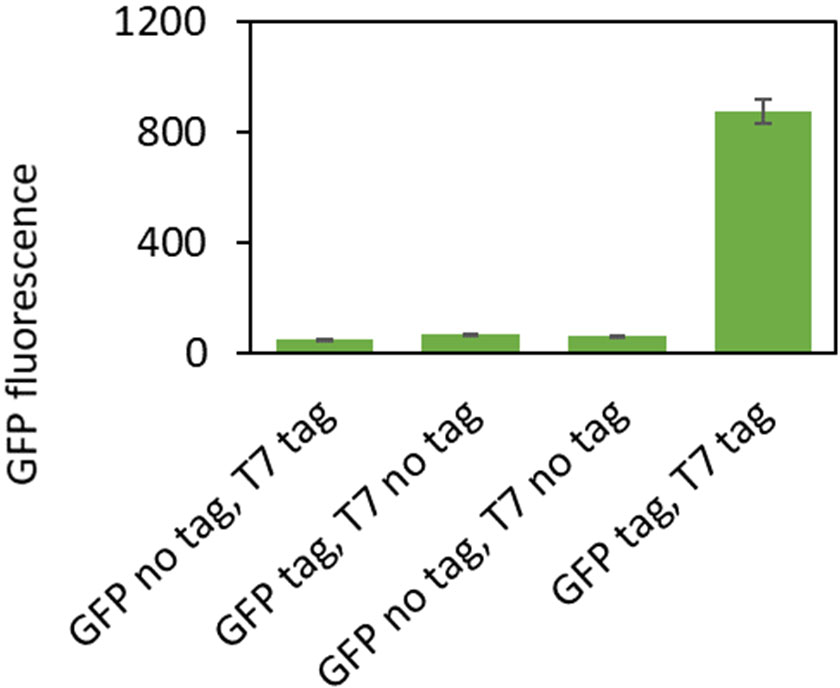
Two synthetic cell populations with or without compatible fusion tags, one containing GFP gene under T7 promoter and the other one containing T7 RNA polymerase, were mixed in equal proportions. The lack of fusion tag is proof of principle control for inoculation experiments shown on Figure 6b.

**Figure S30.**
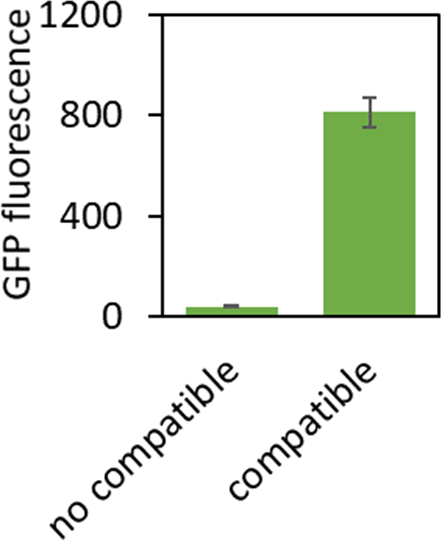
Two synthetic cell populations with compatible or incompatible fusion tags, one containing GFP gene under T7 promoter and the other one containing T7 RNA polymerase, were mixed in equal proportions. The incompatible fusion tag is proof of principle control for inoculation experiments shown on Figure 6b.

**Figure S31.**
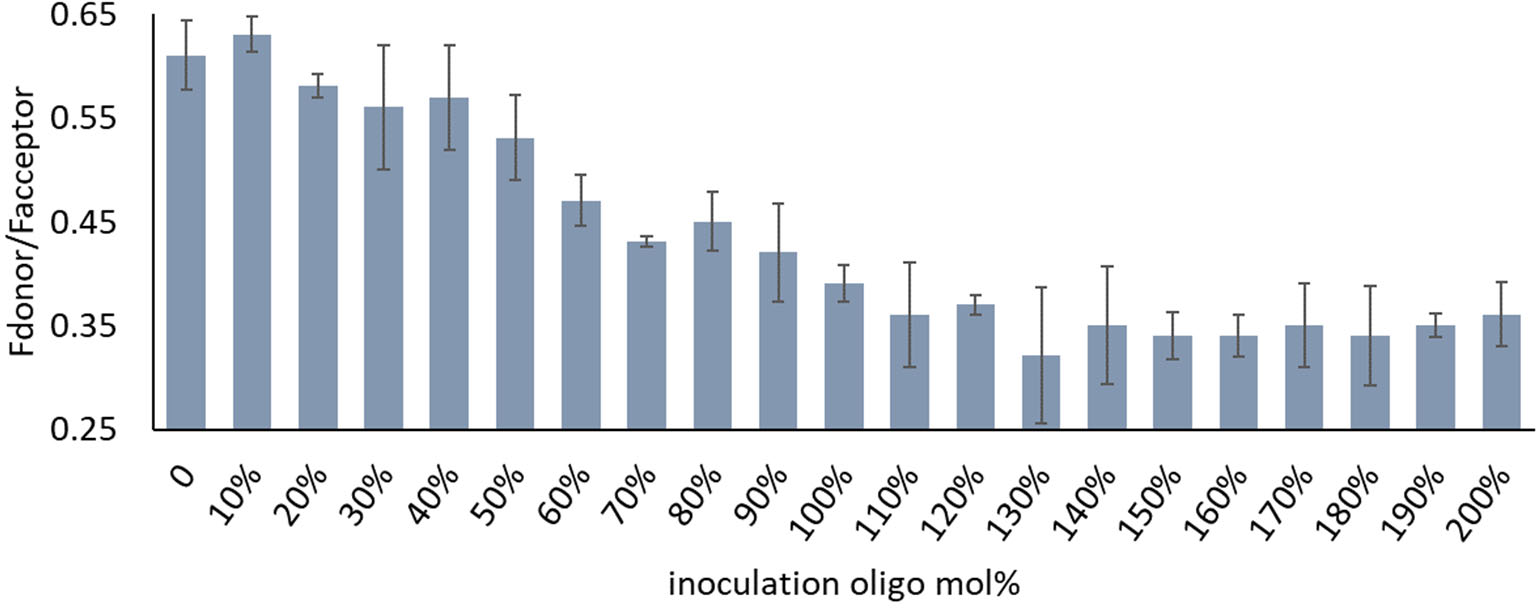
Engineering the inoculation system: oligo complementary to fusion tag was added to vesicles labeled with Fret donor and acceptor dyes at indicted mol% of the total oligo concentration. The calibration curve for Fret experiments is shown on **Figure S34**.

**Figure S32.**
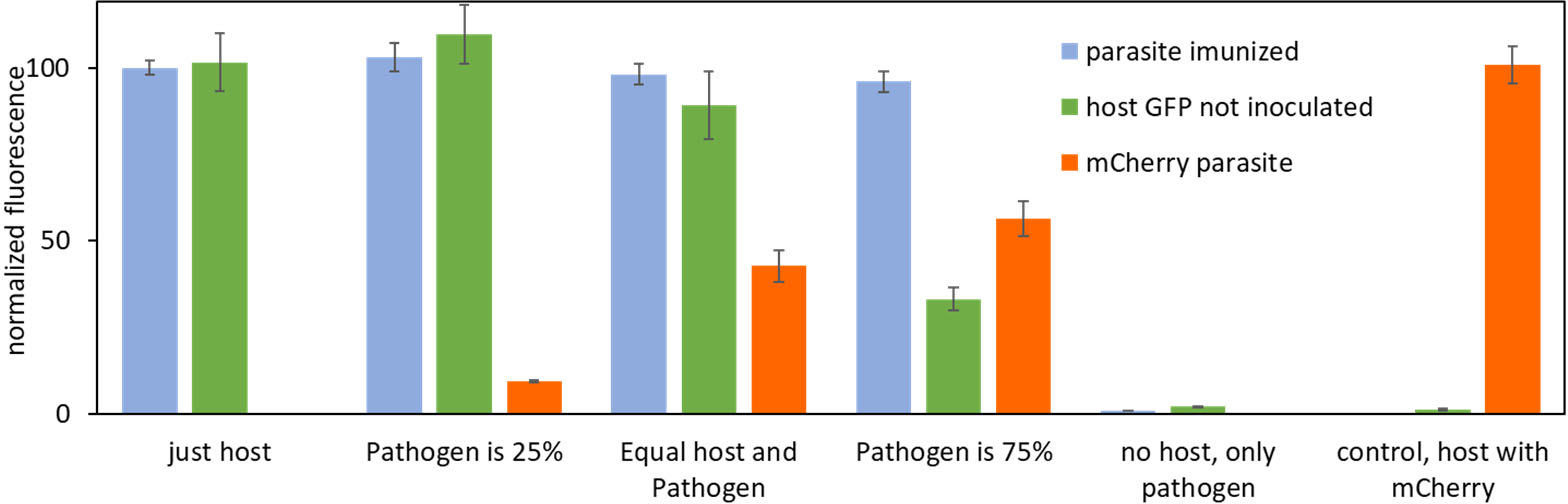
A reversed inoculation was tested by inoculating the parasite instead of host. Error bars indicate SEM, n=3. Fluorescence values are normalized to the blue fluorescence of the membrane dye used to normalize the results to the concentration of host cells.

**Figure S33.**
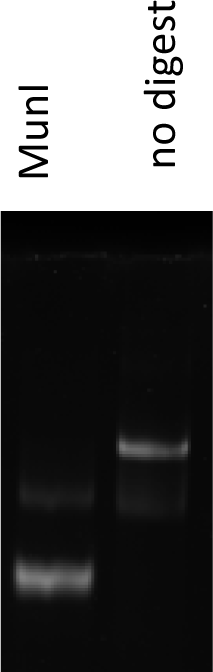
Activity of MunI enzyme expressed in TxTl. A DNA probe was constructed with MunI restriction site and 5’ FAM label. TxTl Expressed MunI from reaction with 5nM plasmid was used in digest experiment, and the product was analyzed on 18% Urea PAGE gel.

**Figure S34.**
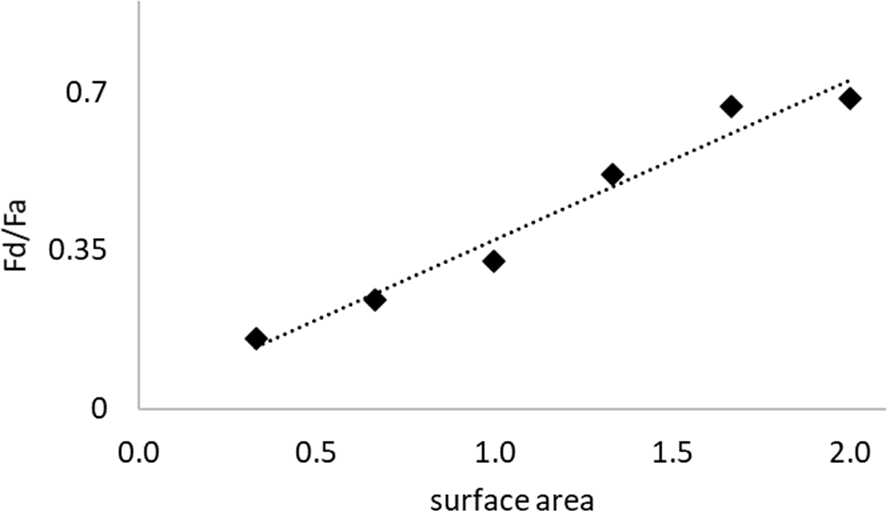
Fret calibration The samples were prepared with varying ratios of the FRET dye pair lipids (Lissamine™ Rhodamine B 1,2-Dihexadecanoyl-sn-Glycero-3-Phosphoethanolamine, Triethylammonium Salt and NBD-PE (N-(7 Nitrobenz-2-Oxa-1,3-Diazol-4-yl)-1,2-Dihexadecanoyl-sn-Glycero-3-Phosphoethanolamine, Triethylammonium Salt)) to the POPC:cholesterol lipid mix, in order to mimic surface area change in fusion experiments. Fd, fluorescence of donor; Fa, fluorescence of acceptor; the relative surface area of 1 is defined as the starting ratio of FRET dyes to lipids in synthetic cell sample.

**Table S1.**
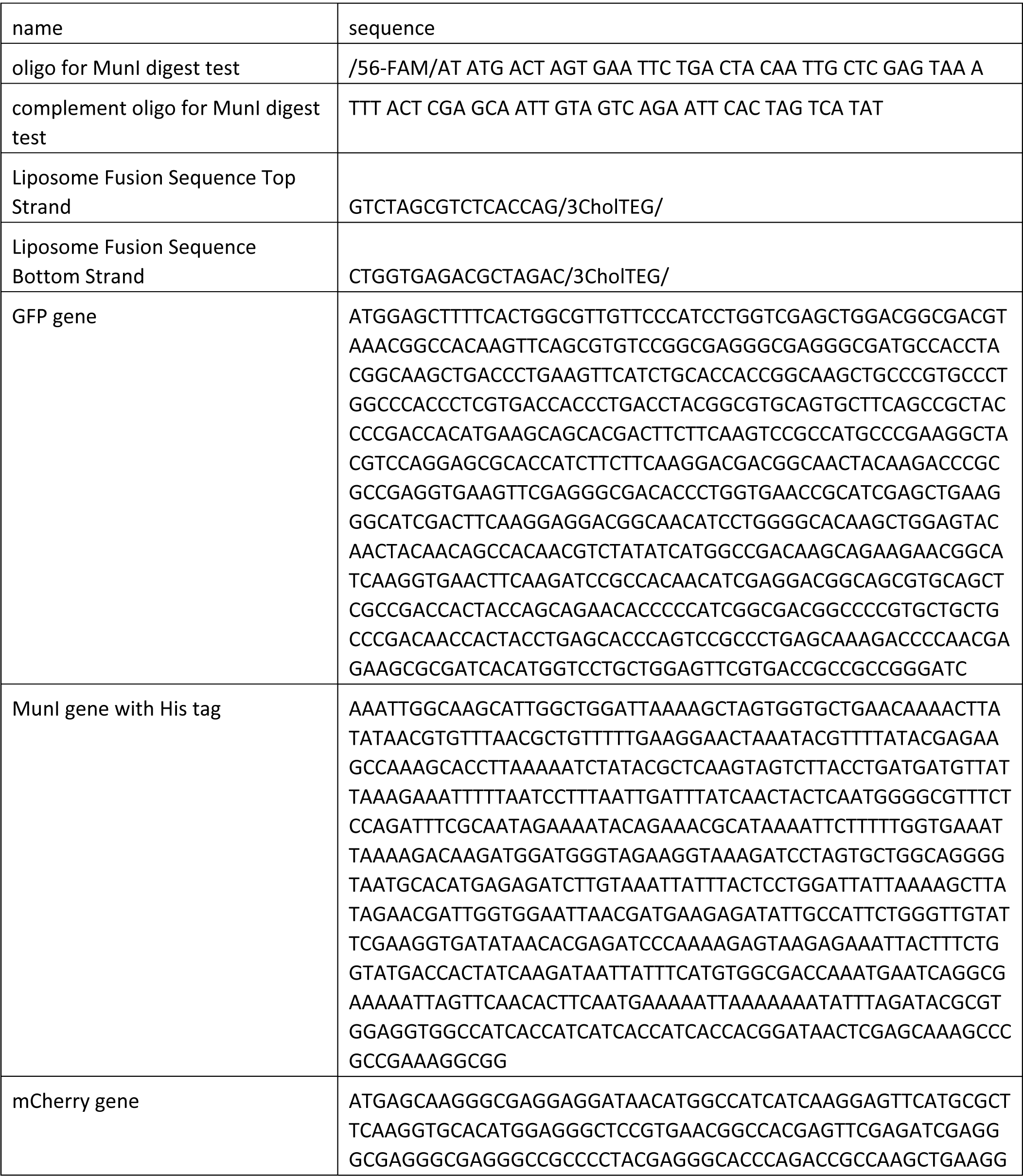
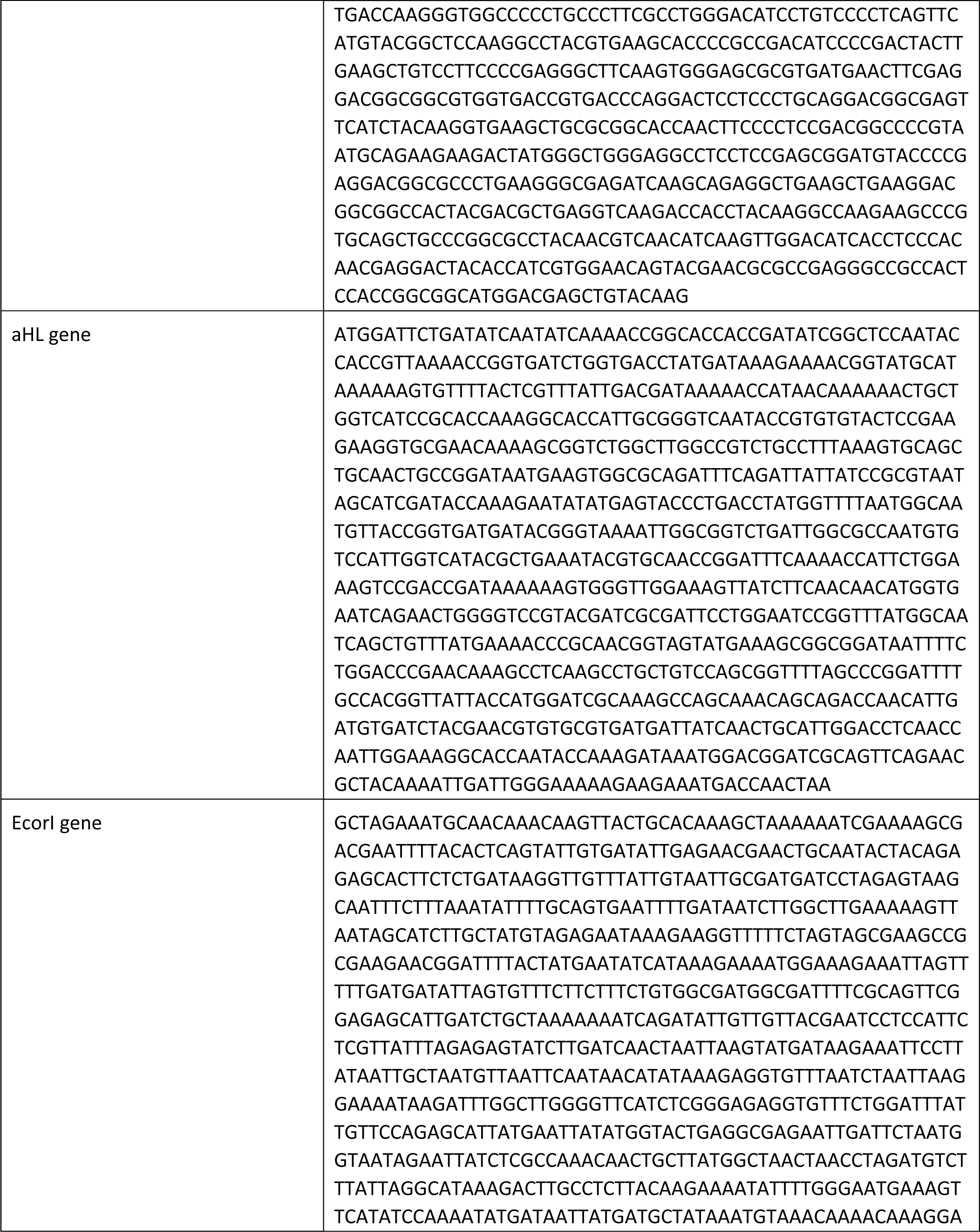
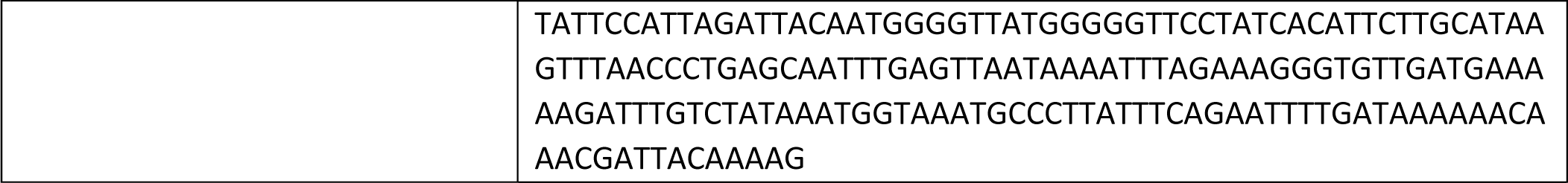
Sequences used in this work.

